# Comprehensive proteomic analysis of JC polyomavirus-infected human astrocytes and their excreted vesicles

**DOI:** 10.1101/2023.04.11.536379

**Authors:** Larise Oberholster, Amandine Mathias, Sylvain Perriot, Emma Blaser, Mathieu Canales, Samuel Jones, Lucas Culebras, Marie Gimenez, G. Campbell Kaynor, Alexey Sapozhnik, Kevin Richetin, Susan Goelz, Renaud Du Pasquier

**Affiliations:** Laboratory of Neuroimmunology, Neuroscience Research Centre, Department of Clinical Neurosciences, Lausanne University Hospital (CHUV) and University of Lausanne, Lausanne, Switzerland; Department of Psychiatry, Center for Psychiatric Neurosciences, Lausanne University Hospital (CHUV) and University of Lausanne, Lausanne, Switzerland; MS & Neurorepair Research Unit, Biogen, Cambridge, MA, USA; Laboratory for Ultrafast Microscopy and Electron Scattering (LUMES), Institute of Physics, Ecole Polytechnique Fédérale de Lausanne (EPFL), Lausanne, Switzerland; Department of Neurology Oregon Health and Sciences University, Portland, OR USA; Service of Neurology, Department of Clinical Neurosciences, Lausanne University Hospital (CHUV) and University of Lausanne, Lausanne, Switzerland

**Keywords:** hiPSC, astrocytes, model, JCPyV, extracellular vesicles, PML

## Abstract

JC polyomavirus (JCPyV) is an opportunistic virus that remains in a latent state in the kidneys of more than half of the human adult population. In rare cases of severe immune suppression, the virus is able to establish a lytic infection of glial cells in the brain, resulting in a debilitating, demyelinating disease known as progressive multifocal leukoencephalopathy (PML). Because of the exceptional species and tissue specificity of the virus, appropriate models of JCPyV infection in the brain are lacking, thus hampering progress towards the development of novel antiviral strategies and biomarkers of disease activity. While PML has traditionally been characterized as a lytic infection of oligodendrocytes, more recent findings suggest an important role for astrocytes during the initial stages of disease. Here, using human induced pluripotent stem cell (hiPSC) derived-astrocytes coupled with a multiparametric approach, we show that 1. JCPyV readily infects and replicates in astrocytes, 2. JCPyV strongly dysregulates the cell biology and 3. these findings adequately reflect *ex vivo* findings. We perform an in-depth characterization of the effect of JCPyV on the cell proteome over time, demonstrating a strong dysregulation of the cell cycle and activation of the DNA damage response. Furthermore, we show that the proteomic signature observed for infected astrocytes is extended to excreted vesicles, underlining their potential to gain valuable insights into JCPyV propagation in the brain.

## Introduction

Progressive multifocal leukoencephalopathy (PML) is a devastating demyelinating disease of the central nervous system (CNS) caused by JC polyomavirus (JCPyV) [1]. JCPyV is an opportunistic virus that normally resides in a benign state in the kidneys and lymphoid organs of more than 50% of the human adult population [2]. However, in rare cases of severe, or selective, immune suppression, the virus is able to reactivate and establish a lytic infection of glial cells in the brain, resulting in severe and irreversible neurological damage. There are still no antiviral strategies against JCPyV and the only means to halt disease progression is to reconstitute an adequate immune response in the brain [3]. The CNS target cells of JCPyV are oligodendrocytes, astrocytes, and to a lesser extent, granule cell neurons of the cerebellum [4–6]. While the lytic infection of oligodendrocytes is what causes demyelination and therefore the hallmark of the disease, the role of astrocyte infection in JCPyV propagation remains unclear. In PML brain lesions JCPyV-infected astrocytes exhibit a bizarre morphology and do not appear to undergo apoptosis [7, 8]. Our limited understanding of JCPyV biology can be attributed to a lack of proper animal models to study the virus in the context of the disease: intracranial inoculation of JCPyV in mice and other species results in tumorigenesis, rather than demyelination, a feature that is not seen in humans [9]. To better characterize JCPyV infection of astrocytes, *in vitro* human glial cell models from diverse origins (primary and transformed glial cells, fetal glial progenitor-derived astrocytes, and induced pluripotent stem cell (hiPSC)-derived glial cells) have been developed [10–15]. Interestingly, findings from both *in vitro* experiments and mice engrafted with human glial progenitor cells suggest astrocytes to enable JCPyV replication earlier than oligodendrocytes, pointing towards an important role for this cell type in the initial stages of PML pathogenesis [16]. Indeed, Kondo *et al.* propose astrocytes and glial progenitor cells (GPCs) to be key perpetuators of JCPyV propagation in the brain and demyelination to be a secondary occurrence. This notion challenges the initial conceptions of PML being primarily a disease of oligodendrocytes and leaves many unanswered questions as to whether the same observations could be expected in the brains of PML patients.

Here, building on previous findings from our laboratory [17], we set out to established a reliable *in vitro* human model of JCPyV infection based on astrocytes differentiated from hiPSCs to: 1.) characterize the effect of JCPyV infection on astrocytes on a cellular level and 2.) perform for the first time an in-depth and comprehensive analysis of the effect of JCPyV on the cell proteome by liquid chromatography-tandem mass spectrometry (LC-MS/MS). Indeed, understanding how the virus influences the host proteome over time is key to defining virus-host interactions. Additionally, we characterize the profile of secreted EVs from JCPyV-infected cells to verify the relevance of using astrocyte-derived EVs found in the periphery as a window into the brain. EVs are membrane-bound structures secreted by all cell types and found in all body fluids as well as in conditioned media from cell cultures [18]. Since EVs reflect the state of their cell-of-origin, they can serve as a rich source of information that can be leveraged for understanding molecular mechanisms at play *in vivo* in an inaccessible organ, such as the brain [19, 20] With this in-depth characterization of JCPyV-infected astrocytes and secreted vesicles, we move closer to defining the role of astrocytes in JCPyV propagation and open new avenues towards the development of antiviral strategies for patients with PML.

## Results

### Establishment of hiPSC-derived astrocytes as an *in vitro* model of early JCPyV infection in the brain

To gain more insights into JCPyV-infected astrocytes in the brain, we developed a new reproducible model based on hiPSC-derived astrocytes from two healthy donors [17]. Cells were infected with 8.6 x 10^3^ Genome Equivalents (GE)/cell and analyzed for their ability to support JCPyV propagation using a multiparametric approach.

Quantitative PCR analysis was performed on the cell pellets and corresponding supernatants of JCPyV-infected and mock-infected cells at 3, 7, 14 and 21 days post-infection (d.p.i.). For the cell pellets, a viral titer of 2.8 x 10^9^ GE/ug cellular DNA was observed at 3 d.p.i. that increased to around 5.2 x 10^11^ GE/ug cellular DNA at 21 d.p.i., indicative of active viral DNA replication (Fig. 1.A). At 3 d.p.i., 7 x 10^7^ GE/ml of culture supernatant was detected that increased rapidly to 3.2 x 10^10^ GE/ml of culture supernatant at 7 d.p.i., suggesting release of virus particles or JCPyV genomic DNA into the culture supernatant. This was then followed by a steady increase until 6.8 x 10^11^ GE/ml of culture supernatant at 21 d.p.i. (Fig. 1.B). We also assessed JCPyV DNA replication on a cellular level by performing fluorescent in situ hybridization (FISH) with JCPyV specific probes on infected and mock-infected cells at 3, 5, 7, 10, 14, 21 d.p.i. (Fig. 1.C,D). JCPyV DNA was restricted to the cell nucleus (Fig. 1.C), with the percentage of cells positive for viral DNA modestly increasing from 3 to 10 d.p.i. (4% - 11%), followed by a statistically significant increase to 35% and 46% at 14 and 21 d.p.i., respectively (Fig. 1.D).

**Figure 1.**
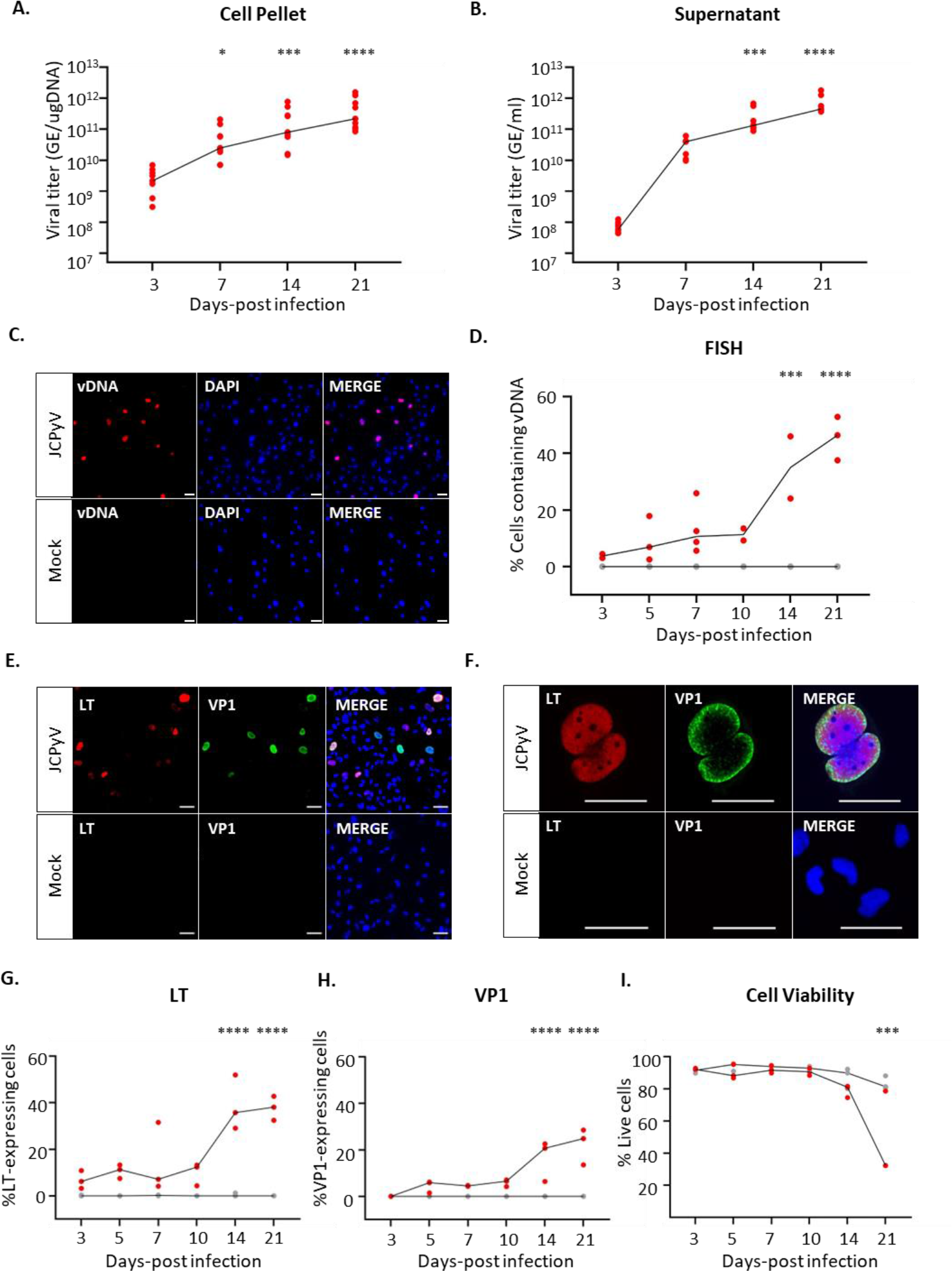
Characterization of JCPyV-infected astrocytes over time. Human iPSC-derived astrocytes were infected with 8.6 x 10^3^ GE/cell JCPyV Mad-1 or mock-infected as a control. At day 3, 5, 7, 10, 14 and 21 cells were analyzed for the presence of viral DNA, expression of viral proteins and cell viability. **A,B.** Quantitative PCR (qPCR) was performed to determine the number of JCPyV DNA copy numbers within the cell pellet (**A**) and culture supernatant (**B**). **C,D.** Fluorescent *in situ* hybridization (FISH) showing cells positive for viral DNA using JCPyV specific probes in infected and mock-infected cells. **C.** Representative images demonstrating JCPyV DNA (red) in the nucleus (DAPI, blue) of hiPSC derived astrocytes at day 7 post-infection (scale bar = 50 µm). **D.** Microscopy quantification of the frequency of cells with viral DNA that co-localized to the cell nucleus (a dot representing the mean of n = 10 images per n = 3 experiments). **E,F.** Representative images with 20x (**E**) and 40x (**F**) magnification confirming the expression of JCPyV early (LT, red) and late (VP1, green) proteins at day 7 post-infection as shown by IFA (scale bar = 50 μm). Cell nuclei were stained with DAPI (blue). **G, H.** Percentage of cells expressing JCPyV LT and/or VP1 was determined by IFA over the course of infection (a dot representing the mean of n = 10 images per n = 3 experiments). **I**. The cell viability was assessed by flow cytometry analysis in JCPyV-infected and mock-infected cells. All panels: n = 3 experiments performed per readout, with each dot on the graph representing an individual experiment (in red: JCPyV; In grey: mock) and the line links the median value of each condition. The effect of infection over time (D3 vs other timepoints; panels D, G, H) and/or as compared to mock-infected conditions (panel I) was tested using a two-way ANOVA followed by Sidak’s multiple comparison test. Statistical significance of data: *p < 0.05; ***p < 0.001; ****p < 0.0001.

The expression of JCPyV large T antigen (LT) and VP1 capsid protein was confirmed in infected astrocytes at day 7 post-infection by immunofluorescence assay (IFA) (Fig. 1.E,F). Cellular staining of LT and VP1 were mostly restricted to cell nuclei that were typically enlarged and comprised a bizarre morphology (Fig. 1.F). The expression kinetics of JCPyV early and late genes were examined by determining the percentage of LT- and VP1-postitive cells at 3, 5, 7, 10, 14, 21 d.p.i. (Fig. 1.G,H). JCPyV LT was detected at 3 d.p.i. in 7% of all quantified cells (Fig. 1.G), whereas VP1 was only detected at the subsequent timepoint (5 d.p.i), comprising 5% of all quantified cells (Fig. 1.H). The percentage of LT- (∼12%) and VP1- (∼5%) expressing cells showed little alterations from 5 d.p.i. to 10 d.p.i. but then increased rapidly to 39% and 17% at day 14 post-infection, respectively. This number then remained constant until 21 d.p.i. when significant cell death (>50%) was observed in the culture as shown by flow cytometry analysis (Fig. 1.I). Cells expressing both the early, LT(+), and late, VP1(+), proteins, comprised only 13% of the culture at the latest timepoint of infection (21 d.p.i.). On the other hand, LT(+)VP1(-) cells made up the biggest percentage of the culture at any timepoint analyzed, representing 22% of all quantified cells at 21 d.p.i. For LT(-)VP1(+) cells, a gradual increase from 0 to 7% from 3 to 21 d.p.i. was observed. All LT(-)VP1(+) cells bore morphologies that resembled dead cells with nuclei that were significantly smaller than that of cells positive for both proteins (results not shown). Since the percentage of LT(+)VP1(-) cells always outnumbered those expressing VP1, it suggests that not all infected cells supported expression of the viral late genes, or that within these cells, the infection was delayed.

JCPyV enters the cell through clathrin-mediated endocytosis whereafter the virus particles are transported to the ER through retrograde trafficking. Once in the ER, JCPyV undergoes conformational changes, allowing its transport to the nucleus where transcription of the virus early genes are initiated [21]. Here, to further characterize the viral particle cellular distribution and the overall cellular state of infected astrocytes, JCPyV-infected and mock-infected cells were analyzed by transmission electron microscopy (TEM) (Fig. 2). Within JCPyV-infected conditions, some cells were found to contain clusters of virions (∼45 nm) associated with the endoplasmic reticulum (ER) cisternae. These cells consisted of intact plasma membranes and cytosols with no obvious indications of JCPyV particles within the nucleus, thereby representing an early stage of the virus life cycle before viral replication (Fig. 2.A). Cells that were at a later stage of infection were identified by densely packed virus particles and tubular structures within the nucleus that coincided with a broken cell membrane and vacuolized cytoplasm (Fig. 2.B). These infected cells underwent a productive viral infection and were identified less frequently than cells showing signs of the early steps of the virus life cycle, i.e., clusters of virions associated with the ER. These findings mirror the results obtained by immune fluorescence analysis and suggests that not all infected cells underwent a productive viral infection, i.e. formation of virus particles or that the infection was significantly delayed.

**Figure 2.**
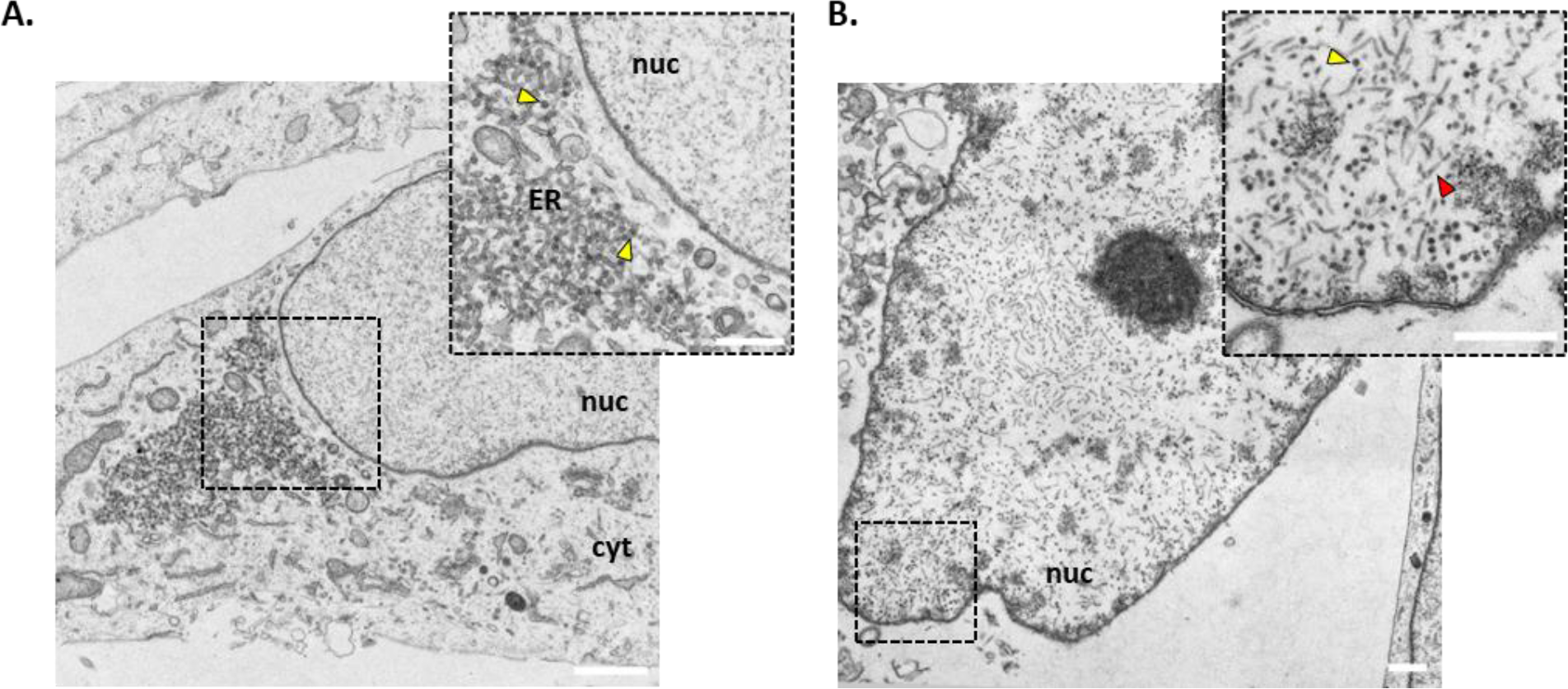
Transmission electron microscopy (TEM) of JCPyV-infected astrocytes. Representative images at day 14 post-infection are depicted confirming the presence of virus particles. **A.** Clusters of JCPyV particles (yellow arrows) were observed within the cytoplasm and found to associate with the endoplasmic reticulum (ER) cisternae. **B.** In some cells, JCPyV particles (yellow arrow) and tubular structures (red arrow) were seen spread throughout the cell nucleus. This coincided with a loss of the cell membrane and cytoplasm, indicative of a later stage of infection (scale bar = 1 µm). See Sup Fig. 1 for the mock-infected control.

### Proteomic signature of JCPyV-infected human astrocytes

Having established our *in vitro* human model of JCPyV-infected astrocytes, we sought to gain deeper insights into the cellular cascades induced upon infection. We therefore set out to perform an in-depth characterization of the cell proteome by liquid chromatography-tandem mass spectrometry (LC-MS/MS) (Fig. 3). Pairwise comparisons were done to identify proteins that were significantly (FDR ≤ 0.05) up- or downregulated in infected conditions as compared to the mock-infected control (Fig. 3.A). As was observed for the viral proteins by IFA, JCPyV LT and small T-antigen (ST) (early gene region) were detected at 3 d.p.i., whereas VP1 and VP2 (late gene region) were only detected at the subsequent timepoint, starting at 7 d.p.i. The expression levels of both ST and LT increased roughly 5-fold from 3 to 21 d.p.i., whereas the expression levels of the viral structural proteins, VP1 and VP2, increased 4-fold and 2-fold, respectively, from 7 to 21 d.p.i. (Sup. Fig. 2).

**Figure 3.**
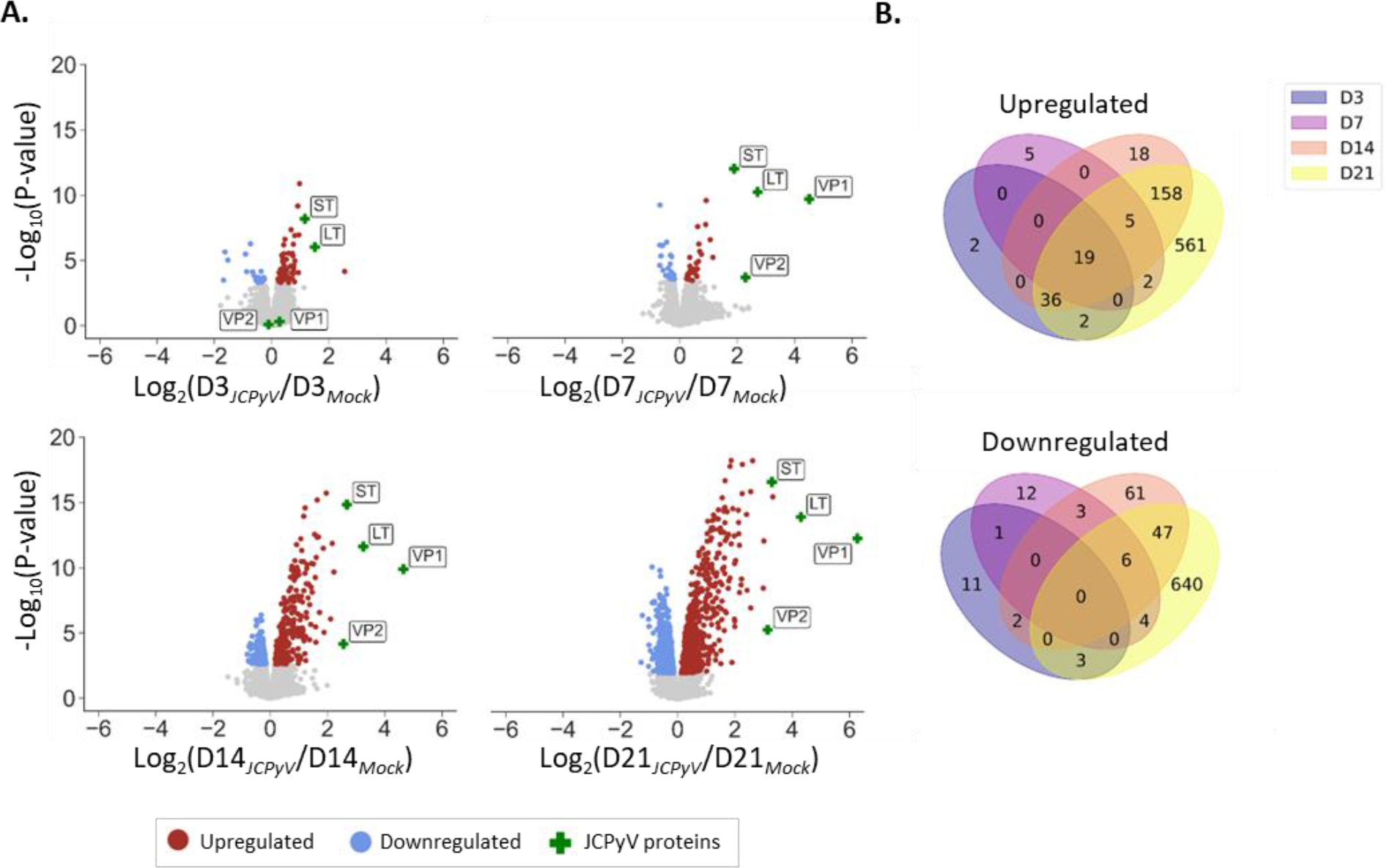
Proteomic analysis of JCPyV-infected astrocytes over time. Cells were infected with JCPyV or mock-infected as described in Fig 1. At day 3, 7, 14 and 21, the cell lysates were collected and analyzed by LS-MS/MS using a TMT labeling approach. **A.** Scatter plots representing the pairwise comparison of quantified proteins in JCPyV-infected cells as compared to the mock-infected control. For each time point (D3, D7, D14, D21) the log2(fold-change) of each quantified protein is represented on the x-axis (JCPyV/Mock) and the corresponding -log10(P-value) on the y-axis. Significantly (FDR ≤ 0.05) dysregulated proteins are shown in color, with upregulated proteins shown in red and downregulated proteins shown in blue. JCPyV proteins are indicated with a green cross. **B.** Venn diagram representing the overlap of significantly up- (red and green, panel A) or downregulated proteins (blue, panel A) at different time points post-infection. Results obtained from n = 3 individual infections.

Regarding the host proteins, among the 7200 proteins detected for all timepoints and conditions analyzed, as many as 783 host proteins were significantly upregulated and 700 proteins significantly downregulated at day 21 post-infection (Fig. 3.B). To gain a better understanding of the types of proteins that were dysregulated by JCPyV, how they behaved over time, and in which cellular pathways they were implicated in, the proteins were scored using the product of |-log10(P-value) * log2(fold-change)| for each protein at the latest timepoint of infection (21 d.p.i.) (Sup. Fig. 3). The hundred highest-ranked proteins, all of which were upregulated, were subjected to protein-protein interaction (PPI) analysis using the STRING database. The network generated consisted of 99 nodes and 1727 edges with an average local clustering coefficient of 0.749, indicative of a highly interactive network. The network was further analyzed using Cytoscape, to include a visual representation of the fold-change (Fig. 4.A). The five proteins with the highest fold-change included: TOP2A; CKAP2, CCNB1; KIF11; BLM, all of which play a role in the mitotic cell cycle process.

**Figure 4.**
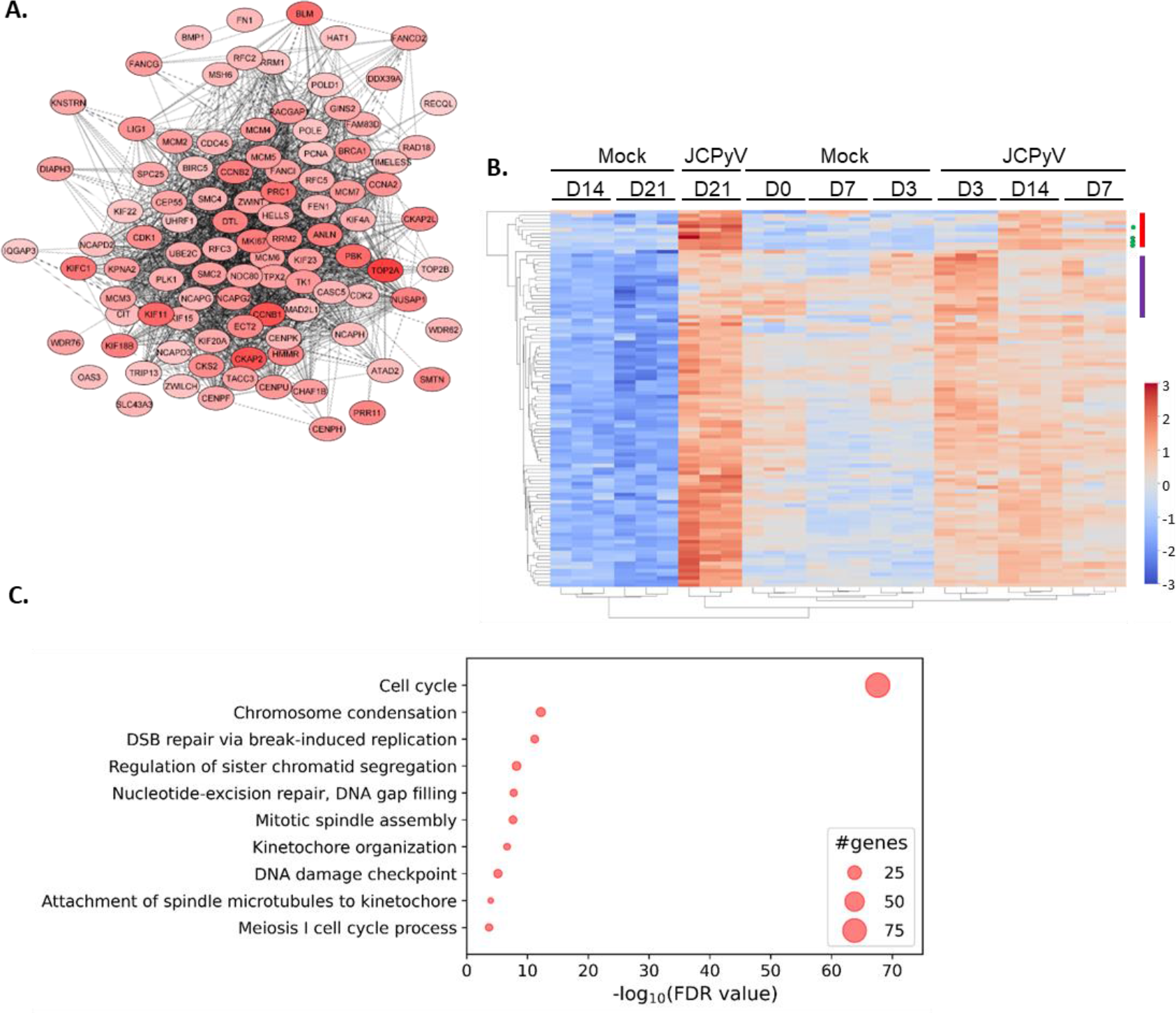
Protein-protein interaction (PPI) analysis and Gene Ontology (GO) enrichment analysis of JCPyV-infected astrocytes. Cells were infected with JCPyV or mock-infected as described in Fig 1. Significantly dysregulated proteins at day 21 post-infection were ranked according to the product of |-log10(P-value) * log2(fold-change)| of which the highest ranking host proteins were selected for further analysis. **A.** Protein-Protein Interaction (PPI) analysis was done on the hundred highest-ranking proteins using the STRING PPI database and visualized on Cytoscape. The color intensity represents the fold-change, the darkest being the most upregulated. **B.** Unsupervised hierarchical Cluster heat map showing the time course of the hundred highest-ranked proteins, where the day post-infection (0, 3, 7, 14 and 21), the condition (JCPyV vs Mock) and the experimental replicates (1-3) are indicated at the top of the image. The viral proteins are indicated with a green dot to the right of the image and notably grouped proteins with coloured bars (see text for details). Results obtained from n = 3 experimental infections. **C.** A bubble chart representing Gene Ontology (GO) enrichment analysis of the hundred highest-ranked proteins. The Biological Processes (BP) are indicated on the y-axis and the corresponding enrichment scores, in terms of the -log10(FDR value), are indicated on the x-axis. The size of the bubble indicates the number of proteins within the corresponding pathway. Results obtained from n = 3 individual infections.

A two-sided unsupervised cluster heatmap was constructed of the hundred highest-ranking proteins to analyze and compare the expression levels of each protein in JCPyV-infected and mock-infected cells over time (Fig. 4.B). The unsupervised heat map highlighted that our samples clustered strongly by culture conditions, e.g. infected vs mock, and time points. First, mock-infected cells at 14 d.p.i and 21 d.p.i clustered further from any other cluster, including early time points of mock-infected cells, suggesting changes in cellular biology as a result of the time spent in culture. This contrasted sharply with the profile of JCPyV-infected astrocytes at 21 d.p.i. that showed a strong upregulation of the same set of proteins that were downregulated in the mock-infected cells, thereby demonstrating the prominent effect of JCPyV on the biology of the cells themselves. Second, mock-infected astrocytes at early time points (0, 3 and 7 d.p.i.) clustered altogether, and away from JCPyV-infected cells at 3, 7 and 14 d.p.i., again highlighting the impact of JCPyV infection on astrocytes even as early as day 3 of infection. When considering the intensity of expression of the proteins in the heat map, in addition to the general tendency of most of the proteins to be gradually upregulated by JCPyV infection, we were able to recognize specific patterns of expression. A first set of proteins with the highest Z-scores at 21 d.p.i. in JCPyV-infected conditions, including viral proteins, LT, ST, VP1 and VP2 (Fig. 4.B., green dot) and host proteins FANCG; CENPU, NCAPD3; BMP1; RFC3 and TOP2B, were shown to evolve together (Fig. 4.B, red bar). The level of expression of these proteins remained relatively stable in all mock-infected conditions over the entire time course, however, the level of these proteins gradually increased in the JCPyV conditions, pointing towards a group of proteins upregulated by JCPyV but not affected by the culture time. Interestingly, a second set of proteins (Fig. 4.B, purple bar: KIF23; PLK1; NDC80; CEP55; CKAP2L; NUSAP1; KIF15; IQGAP3; PCNA; KNSTRN; NCAPH; GINS2; ZWILCH; TACC3; RAD18:

UBE2C; UHRF1) were found with lower Z-scores in JCPyV-infected astrocytes at 21 d.p.i. than at 3 d.p.i., suggesting they might play an important role during the early stages of infection.

To identify cellular pathways that were perturbed by JCPyV, we performed Gene Ontology (GO) enrichment analysis of the hundred highest-ranked proteins (Fig. 4.C). As such, we were able to define an important enrichment of proteins involved in the cell cycle, which comprised 79% of the hundred highest-ranking proteins and thereby showed a high degree of redundancy with other GO terms (Sup. Table 1). Proteins associated with the S and G2/M phases of the cell cycle were especially enriched, including cyclin proteins, CCNA2, CCNB1, CCNB2 and cyclin-dependent kinases, CDK1 and CDK2. Other enriched GO terms included those involved in mitotic processes, such as chromosome condensation, regulation of sister chromatid segregation, mitotic spindle assembly, kinetochore organization, attachment of spindle microtubules to kinetochores and the meiosis I cell cycle process. Taken together, these data suggest that JCPyV facilitates cell cycle progression and arrest at the G2/M phase as has been extensively shown for JCPyV [11, 22–24] and other polyomaviruses such as BK virus [25] and SV40 [26]. It was reported that JCPyV-infected human neuroblastoma IMR-32 cells accumulate in the G2 phase of the cell cycle by activating ATM- and ATR-mediated checkpoint pathways that form part of the cell’s DNA damage response (DDR) [24]. Interestingly, from our own data, the three GO terms: DNA damage checkpoints, nucleotide excision repair via DNA gap filling and double-strand break (DSB) repair via break-induced replication were also among the top 10 most significantly enriched pathways in JCPyV conditions and represented 23% of the hundred most significantly upregulated proteins in JCPyV-infected astrocytes.

**Table 1.** Antibody list used for Immunofluorescence assay

| Target | Clone | Host | Dilution | Manufacturer | Reference |
| --- | --- | --- | --- | --- | --- |
| SV40 T-antigen | EPR22694-148 | Rabbit | 1/200 | Abcam | ab234426 |
| VP1 | PAB597 | Mouse | 1/800 | Biogen |  |
| Phospho-ATM (Ser1981) | 10H11.E12 | Mouse | 1/100 | Sigma-Aldrich | 05-740 |
| H2AX (pS139) | N1-431 | Mouse | 1/100 | BD Biosciences | 560443 |
| Phospho-Chk2 (T68) | 1238H | Rabbit | 1/100 | R&D systems | MAB1626-SP |
| BRCA1 (Ab-1) | Ms110 | Mouse | 1/100 | Sigma-Aldrich | OP92 |
| p53BP1 (Ab-1) | pAb | Rabbit | 1/100 | Sigma-Aldrich | PC712 |
| FANCD2 | pAb | Rabbit | 1/200 | GeneTex | GTX30142 |
| PLK | F-8 | Mouse | 1/100 | Santa Cruz<br>Biotechnology | sc-17783 |

Since all polyomaviruses, including JCPyV, rely on the DDR pathways for productive viral infection, we next wanted to confirm the activation of these pathways on a cellular level in infected astrocytes [25]. Once DNA damage occurs, a cellular cascade is activated that allows for the recruitment of multiple repair factors to the site of the lesion. This includes the DNA damage checkpoint protein, ataxia telangiectasia mutated (ATM) that, once activated, is able to phosphorylate histone variant H2AX at serine 139 (yH2AX) and protein kinase CHEK2 that serve as a signal for the recruitment of DNA repair proteins, including 53BPI, BRCA1 and FANCD2, resulting in the formation of nuclear foci [27, 28]. From our proteomic data, we confirmed a temporal increase in the relative abundance of proteins involved in the aforementioned processes that mostly reached significance from 14 d.p.i. and onward (Fig. 5.A). The formation of nuclear foci (yH2AX; p53BPI; FANCD2; BRCA1) and the upregulation of DNA damage checkpoint proteins (ATM, CHEK2, PLK) were then further confirmed on a cellular level by IFA in JCPyV-infected cells (Fig.5.B) but were not observed in the mock-infected control (Sup. Fig.4). This indicates that in our astrocytes, JCPyV induces the activation of DDR pathways similarly to what has been reported in primary human astrocytes infected with JCPyV and PML brain lesions [23].

**Figure 5.**
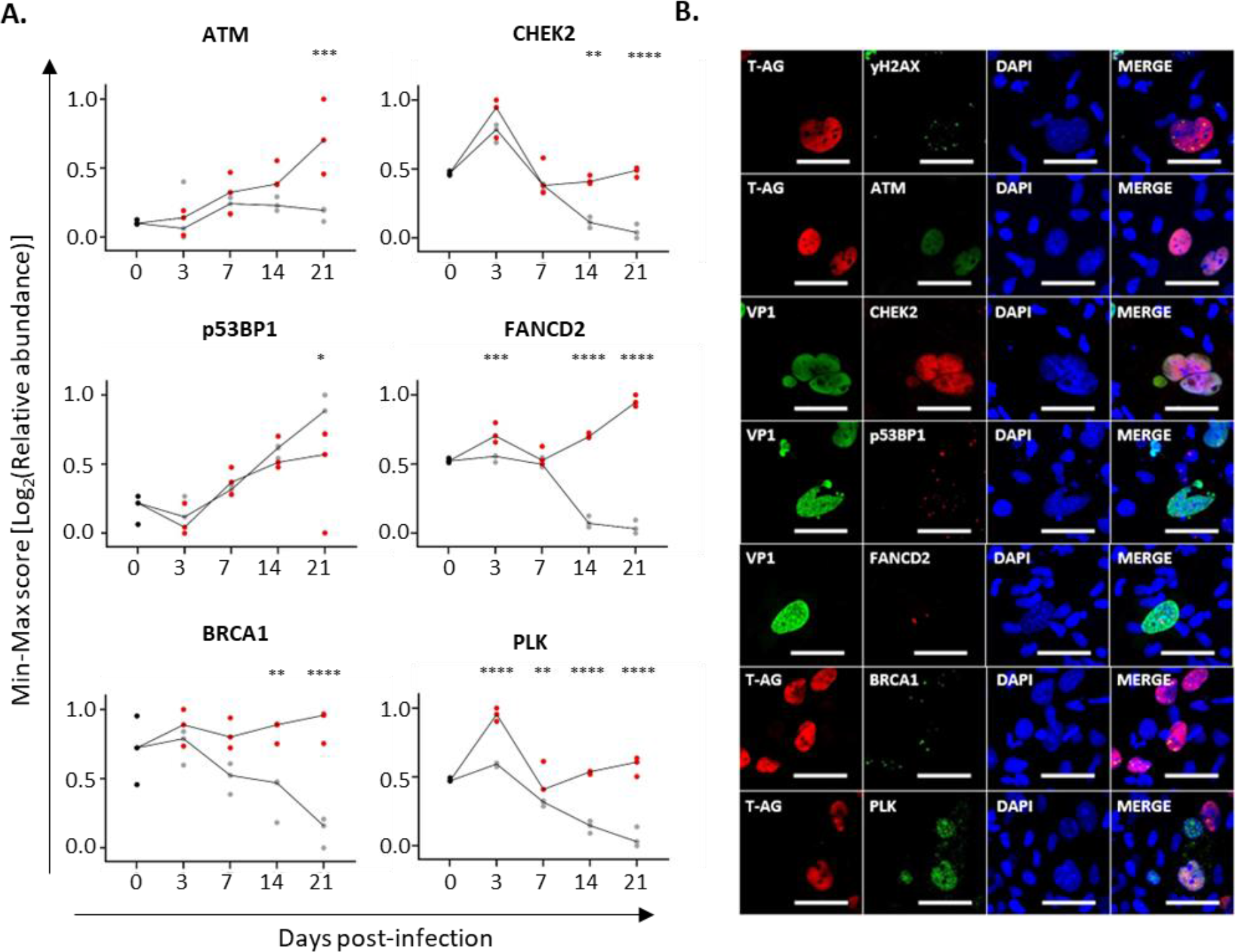
Activation of the DNA damage response (DDR) in JCPyV-infected astrocytes. **A.** Min-Max score of the log2(Relative abundance) of DDR proteins quantified over time by proteomic analyses in JCPyV-infected (red) and mock-infected (grey) astrocytes. A minimum of n = 3 experiments were performed, with each dot on the graph representing an individual infection and the line that links the median value of each condition (JCPyV vs Mock). The asterisks (*) represent significant differences between infected and mock conditions for each protein using a two-way ANOVA followed by Sidak’s multiple comparison test. Statistical significance of data: *p < 0.05; **p<0.01; ***p < 0.001; ****p < 0.0001. **B.** Representative images taken at 7 d.p.i showing the activation of the DDR through upregulation of DNA damage checkpoint proteins (ATM, CHEK2, PLK) and the formation of nuclear foci (yH2AX, p53BP1, BRCA1, FANCD2) in infected astrocytes expressing either JCPyV LT (red) or VP1 (green) (scale bar = 50 µm). See Sup Fig. 4 for the mock-infected control.

To conclude, this in-depth characterization of the effects of JCPyV on the proteome of astrocytes confirms the current knowledge on the biology of JCPyV and other polyomaviruses, i.e., host-pathogen interactions involved in the cell cycle and DDR pathways. Since our human *in vitro* hiPSC-derived model appeared to encapsulate what has been observed for the virus both in primary human astrocytes in culture and in PML brain lesions, we used it to further explore JCPyV biology in astrocytes and its effects on EVs released from infected cells.

### Proteomic analysis of EVs from JCPyV-infected astrocytes

The ability of brain-derived EVs to cross the blood-brain-barrier (BBB) has piqued the interest of neuroscientists for their potential to serve as rich sources of information pertaining to inaccessible organs, such as the brain, particularly in the context of disease [20]. Having demonstrated that the proteomic signature of our *in vitro* model of JCPyV infection in the brain accurately reflects features observed for infected astrocytes *in vivo*, supernatants were collected from JCPyV-infected and mock-infected astrocytes at day 7 and 14 post-infection. These time points were selected to avoid significant cell death (refer to Fig. 1.I), in the culture that would risk contamination of the EV preparations. EVs were isolated using the classical method of differential ultracentrifugation, whereafter the EV-enriched fractions were analyzed by LC-MS/MS (Fig. 6.A). To confirm that EVs from JCPyV-infected astrocytes constituted a proteomic signature specific to viral infection and distinct from inflammatory conditions, EVs were also collected from astrocytes stimulated with proinflammatory cytokines, TNF-α and IL-1b, as well as unstimulated, resting controls. These two cytokines have been shown to be key players in neuroinflammatory diseases [17, 29]. To assess the purity of the EV-enriched fractions, the relative abundances of proteins normally found to associate with EVs were compared to that of potential contaminants, according to MISEV2018 specifications, for each sample [30]. (Sup. Fig. 5). EV-associated proteins (from categories 1A, 1B, 2A, 2B) were significantly enriched across all samples analyzed as compared to contaminant proteins (from categories 3A, 3B), suggesting that a good level of purity was reached. Notably, EVs generated under infected conditions contained a high abundance of ribosomal proteins from category 3B as compared to EV generated under stimulated or resting conditions. Since all EV isolations were done using the same procedure, we suspected the association of ribosomal proteins with EVs to be specific to viral infection.

**Figure 6:**
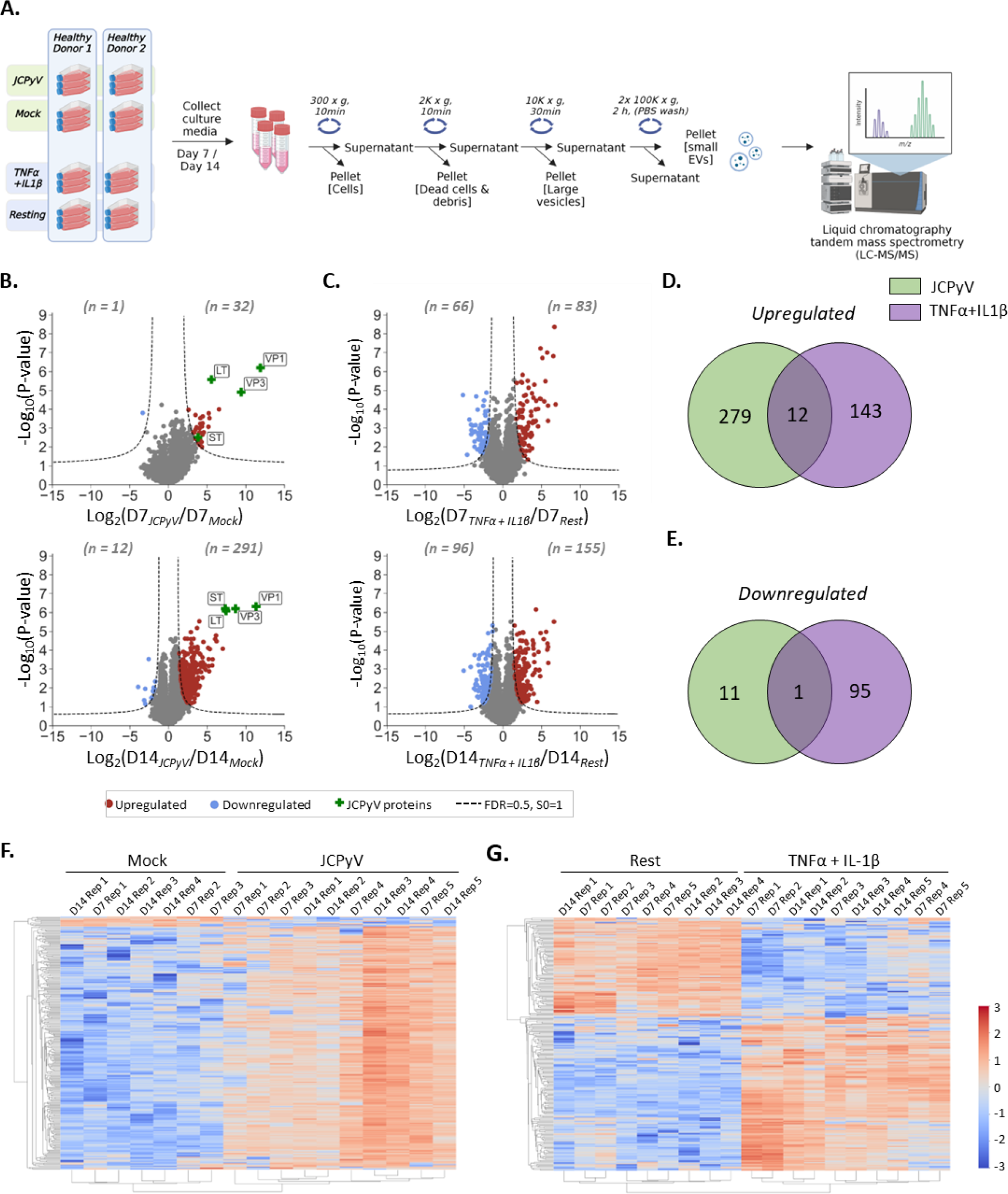
Proteomic signature of extracellular vesicles (EVs) from JCPyV-infected or cytokine-stimulated astrocytes. Human iPSC-derived astrocytes were infected with 8.6 x 10^3^ GE/cell JCPyV or stimulated with 10ng/ml TNFα and 10ng/ml IL-1β. As the negative control for each condition, the cells were either mock-infected or left resting, respectively. Results were obtained from n = 4-5 independent experiments. **A.** At day 7 and 14, EVs were collected from the culture supernatant of the different conditions and subjected to increasing speeds of centrifugation to remove cells, dead cells, debris and large vesicles, e.g. microvesicles (MVs) and apoptotic bodies. The final pellet from the 100K x g centrifugation step, comprising small EVs, was collected and analyzed by liquid chromatography tandem mass spectrometry (LS-MS/MS) using a label-free quantification method. **B.** Scatter plots showing quantified proteins in JCPyV conditions as compared to mock-infected control. For each timepoint (D7 and D14) the Log2(fold-change) of the quantified proteins are represented on the x-axis (JCPyV/ Mock) and the corresponding -Log10(P-value) on the y-axis. Significantly (FDR ≤ 0.05) dysregulated proteins are shown in color, with upregulated proteins shown in red and downregulated proteins shown in blue. JCPyV proteins are indicated with a green cross. **C.** Scatter plots showing quantified proteins from TNFα + IL-1β -stimulated astrocytes as compared to resting (i.e. unstimulated) astrocytes, similarly as described in B. Upregulated proteins are shown in red and downregulated proteins in blue. **D, E.** Venn diagrams showing the overlap of significantly upregulated (**D**) or downregulated (**E**) proteins in JCPyV-infected or TNFα+IL-1β-stimulated astrocytes at day 14 post-infection or post-stimulation, respectively. **F, G.** Unsupervised cluster heatmaps showing the time course of the most significantly dysregulated proteins (taken from day 14) in JCPyV-infected (**F**) or TNFα+IL-1β-stimulated (**G**) conditions as compared to the mock-infected or resting controls, respectively.

Pairwise comparisons of the EV-enriched fraction proteomic data were done for all time points to identify proteins that were significantly (FDR ≤ 0.05; S0 = 1) dysregulated in infected (Fig. 6.B) or cytokines-stimulated (Fig. 6.C) conditions as compared to the mock-infected or resting controls, respectively. In infected conditions, JCPyV early (LT and ST) and late (VP1 and VP3) proteins were detected in the EV-enriched samples of all the time points analyzed. We cannot rule out that there was an overlapping size distribution of JCPyV particles (∼45 nm) and small EVs (30-150 nm), resulting in their co-sedimentation following high-speed ultracentrifugation. Of main interest, however, was the remarkable dysregulation of host proteins induced by the virus. The number of significantly upregulated (n = 291) proteins starkly outnumbered those that were significantly downregulated (n = 12) at day 14 post-infection and sharply contrasted with the proteins identified in cytokine-stimulated conditions at both timepoints analyzed. Of the 291 significantly upregulated proteins in JCPyV conditions, only 12 proteins (4%) were shown to overlap with the 155 significantly upregulated proteins in cytokine-stimulated conditions (Fig. 6.D). These proteins included: ILF3; SRSF1; U2AF2; ISG15; DDX17; HNRNPAB; ELAVL1; HNRNPL; HNRNPM; HNRNPH3; TM9SF4 and DDX5 that mostly have functions associated with mRNA splicing. Correspondingly, of the 12 significantly downregulated proteins at 14 d.p.i. in JCPyV conditions, only SLC38A3 formed part of the 96 significantly downregulated proteins in cytokine-stimulated conditions (Fig. 6.E). We next constructed an unsupervised cluster heat map to compare the relative abundances of significantly dysregulated proteins (taken from day 14) over time for JCPyV-infected or cytokine-stimulated conditions as compared to the mock-infected or resting control, respectively (Fig. 6.F, G). JCPyV conditions clustered separately from that of the mock-infected control, with no specific clustering based on the time point analyzed (Fig. 6.F). This suggested that the time course from 7 to 14 d.p.i. had little to no effect on the protein levels. In a similar way, column-clustering resulted in the separate grouping of cytokine-stimulated conditions from the resting controls and showed no apparent effect of time (Fig. 6.G).

To gain a better understanding of the cellular pathways in which these proteins were implicated, we performed PPI analysis and GO enrichment analysis of the hundred most dysregulated proteins from each condition according to the product of |-log10(P-value) * log2(fold-change)| (Sup. Fig. 6). For infected conditions, the generated STRING network consisted of 100 nodes and 500 edges with an average local clustering coefficient of 0.502, whereas the network for cytokine-stimulated conditions consisted of 99 nodes and 194 edges with an average local cluster coefficient of 0.479. Both networks were further analyzed using Cytoscape to incorporate visualization of the fold-change. For JCPyV conditions, the five proteins with the highest fold-change included: TOP2A, SMC4, MCM3, KIF11 and FEN1 thereby comprising a proteomic signature highly representative of what was observed for the virus on a cellular level (Fig. 7.A). DNA topoisomerase II alpha (TOP2A) was shown to be the most highly upregulated host protein in both the JCPyV-infected cells and the EV-enriched fraction, with a fold-change of 5 and 130, respectively at 14 d.p.i. Interestingly, the same protein was mostly absent in the EV fraction generated from astrocytes using the other conditions tested (mock-infected, cytokine-stimulated, resting). The proteomic signature of the EV-enriched fraction from cytokine-stimulated conditions was starkly different from that of the EV-enriched fraction generated under infection conditions. Under cytokines-stimulated conditions, the top five proteins with the highest fold-change in the EV-enriched fractions included: MX1, GGT5, BST2, DDX58 and IFIT1. Intriguingly, besides GGT5, these proteins all play a role in the defense response against viruses and were completely absent from the EV fraction generated under infection conditions (Fig. 7.B).

**Figure 7:**
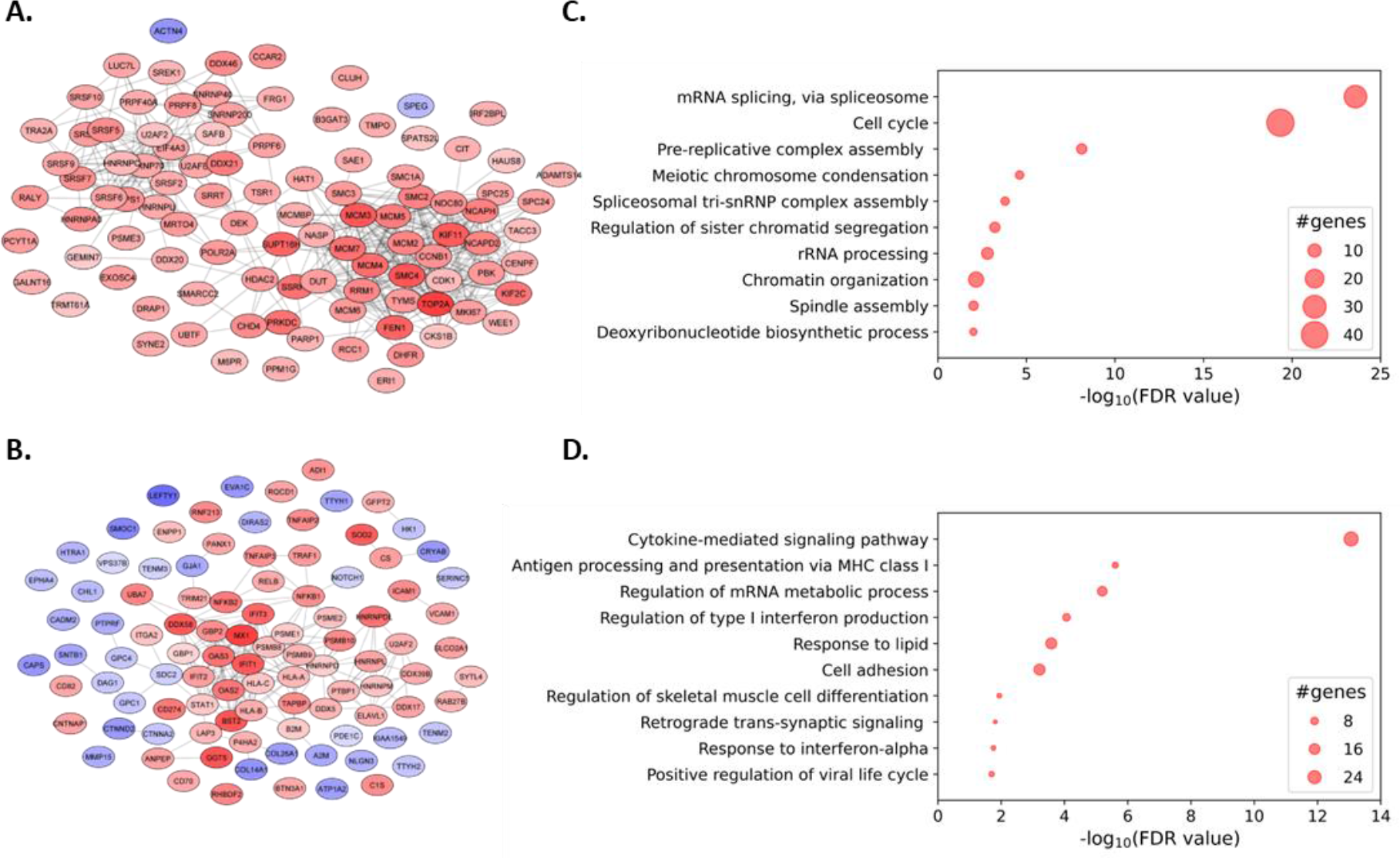
Protein-protein interaction (PPI) analysis and Gene Ontology (GO) enrichment analysis of EVs from JCPyV-infected or TNFα+IL-1β-stimulated astrocytes. Significantly dysregulated proteins at day 14 of infection or stimulation were ranked according to the product of |-log10(P-value) * log2(fold-change)| of which the highest-ranked host proteins were selected analysis. **A, B.** PPI networks generated using the STRING database for the JCPyV (**A**) and TNFα+IL-1β (**B**) conditions were further analyzed using Cytoscape. Upregulated proteins are shown in red and downregulated proteins in blue, with the fold-change represented by the color intensity, the darkest being the most dysregulated. **C, D.** GO enrichment analysis was performed on the hundred highest-ranking proteins of JCPyV (**C**) and TNFα+IL-1β-stimulated (**D**) conditions. The Biological Processes (BP) are indicated on the y-axis and the corresponding enrichment scores, in terms of the -log10(FDR value), are indicated on the x-axis. The size of the bubble correlates with the number of genes within the corresponding pathway.

We next performed GO enrichment analysis on the same set of top-ranked proteins from either JCPyV-infected (Sup. Table. 2) or cytokine-stimulated (Sup. Table. 3) conditions as compared to the mock-infected or resting control, respectively. In the EV-enriched fraction from JCPyV infected cells, proteins involved in mRNA splicing via spliceosomes and the cell cycle were highly enriched and to a lesser extent, proteins involved in pre-replicative complex assembly and mitotic processes, e.g. chromosome condensation, regulation of sister chromatid segregation and chromatin organization, thereby reflecting what was observed for the virus on a cellular level (Fig. 7.C). When considering all the significantly upregulated proteins in the EV-enriched fraction from JCPyV-infected cells, roughly 25% were shared with those significantly upregulated in infected astrocytes. These proteins were primarily implicated in the cell cycle, whereas the remaining 75% comprised in the EV-enriched fraction only was primarily implicated in RNA processing. Once again, these observations were highly different from that of the EV-enriched fractions from cytokine-stimulated cells, of which the five GO terms with the highest scores included cytokine-mediated signaling pathways, followed by antigen processing and presentation via MHC class I molecules, regulation of mRNA metabolic processes, regulation of type I interferon production and response to lipids (Fig. 7.D). Taken together, these data show that EVs from JCPyV-infected astrocytes have a very distinct proteomic signature from those generated under inflammatory conditions. The proteomic content of these EV-enriched fractions somewhat reflects the dysregulation brought on by the virus on a cellular level, but also indicates selective packaging of some proteins into EVs. This feature opens interesting avenues regarding cellular communication in the context of JCPyV infection and highlights the potential of EVs as a tool to gain better insights into JCPyV biology in the brain of PML patients.

## Discussion

In this study, we first set out to establish a reliable human *in vitro* model of JCPyV infection in astrocytes. Using a variety of techniques, we show that hiPSC-derived astrocytes support JCPyV infection similarly to primary human astrocytes [22, 23]. This was demonstrated by a support of viral DNA replication and expression of the virus early and late genes. Temporal analysis revealed a small percentage of cells positive for JCPyV DNA or viral proteins during the first ten days of infection that rapidly increased during the later stages of infection, e.g. 14 d.p.i and onwards. It has been reported that the JCPyV infectious cycle in primary human astrocytes is delayed as compared to that of SVG-A cells in which the presence of SV40 LT accelerates the production of the viral late genes [22]. While SVG-A cells are widely used to study JCPyV biology [31], the constitutive expression of SV40 LT represents a barrier to bridge *in vitro* findings to what can be expected *in vivo*. Conversely, we found that the number of LT-expressing cells always outnumbered those expressing VP1, even during the later stages of infection. Additionally, TEM analysis of JCPyV-infected astrocytes revealed a higher frequency of infected cells devoid of signs of JCPyV replication (virion clusters associated with the ER but absent in the nucleus) as compared to newly produced JCPyV virions (virus assembly within the nucleus). Similar observations were made in PML brain lesions [6]. These findings suggest that not all infected astrocytes supported a productive viral infection or that in some infected cells, the expression of the virus late genes, and formation of virus particles, is delayed. A productive viral infection was discernible in cells, which we suspected to be dead or dying, having nuclei filled with virus particles and tubular structures that coincided with a broken plasma membrane and vacuolized cytoplasm. Supporting these findings, Seth *et al*. found that human glial progenitor derived astrocytes that were permissively infected with JCPyV eventually led to non-apoptotic cell death that bore striking similarities to necrosis, e.g vacuolized cytoplasm and dissolution of organelles [12]. Mazlo *et al.* made similar observations in PML brain biopsies [6]. As such, our model appears to faithfully recapitulates not only features of JCPyV infection *in vitro* but all characteristics known for the infected astrocytes in PML brain lesions. To further our investigation into the biology of JCPyV-infected astrocytes, we performed an in-depth characterization of the cell proteome by liquid chromatography tandem mass spectrometry LS-MS/MS. We found that JCPyV had a remarkable impact on the biology of astrocytes, resulting in a high number of significantly dysregulated proteins. Notably, proteins involved in the cell cycle and cell division were significantly upregulated in the JCPyV condition and the level of these proteins correlated with the time course of infection. Proteins involved in the G2/M phase of the cell cycle were especially enriched, suggesting that the virus was able to facilitate cell cycle progression as has been reported for JCPyV [11, 22] and other polyomaviruses [26, 32]. Additionally, proteins involved in the DNA damage response (DDR) were also highly enriched. JCPyV LT was shown by others to mediate cell cycle arrest at the G2/M phase in a human neuroblastoma cell line through the activation of ATR and ATM-mediated pathways [24]. Furthermore, DDR proteins have previously been reported to be upregulated in primary human astrocytes infected with JCPyV *in vitro* and bizarre astrocytes from PML brain lesions [23]. In a more recent study, DDR proteins were described to be localized to sites of JCPyV DNA replication within the nuclei of primary human astrocytes [33]. The involvement of the DDR in the propagation of other polyomaviruses, including BK virus [25], Merkel cell polyomavirus (MCPyV) [34], SV40 [35] and murine polyomavirus (MPyV) [36] has also been well characterized. Here, our data further confirm that proteins involved in DNA damage checkpoints, including ATM, CHEK1, CHEK2, BRCA1, FANCD2 and PLK, are upregulated in JCPyV infected cells and form nuclear foci, which along with the phosphorylation of H2AX, indicate activation of DDR pathways. Overall, our systematic analysis of JCPyV-infected astrocytes not only accurately depicts features of the virus life cycle, as discussed above, but also the altered biological state observed for JCPyV-infected primary human astrocytes *in vitro* and *ex vivo*, thus highlighting the relevance of our model to gain further insights into JCPyV pathophysiology in the brain [11, 23, 24, 33]. In addition to characterizing the effect of JCPyV on the cell proteome, we wanted to establish whether the same signature was extended to EVs from infected astrocytes. In the absence of proper animal models to study JCPyV in the context of PML, brain-derived EVs found in the blood provides the next best solution to bridge *in vitro* findings to what can be expected *in vivo*. Given that our model proved to be highly reproducible and representative of what could be expected for infected astrocytes in PML brain lesions, we investigated whether astrocyte-derived EVs would reflect the state of infection and provide insights into the cellular mechanisms at play. Towards this aim, EVs were isolated from JCPyV-infected and mock-infected astrocytes and analyzed by LC-MS/MS. First, JCPyV viral proteins (LT, ST, VP1 and VP3) were found to co-isolate with EVs. While this might be attributed to the co-sedimentation of JCPyV particles and EVs following high-speed centrifugation, we do not rule out the possibility that JCPyV can propagate through EVs. Indeed, EV-mediated transfer of JCPyV has been reported in SVG-A [37, 38] and choroid plexus epithelial cell cultures [39] and *ex vivo* through the association of JCPyV VP1 and genomic DNA with plasma derived EVs [40]. Similar to the observations made in our hiPSC-derived astrocyte model, we found that JCPyV had a prominent impact on the proteomic content of the EV-enriched fraction from these infected cells. Proteins involved in cellular processes such as mRNA splicing via spliceosomes, cell cycle and mitotic processes were particularly enriched, thereby in part reflecting what was observed for the virus in JCPyV-infected cells. Importantly, only about 25% of the proteins from the EV fraction was shared between the cellular fraction and the EV-enriched fraction. The remaining 75% delineated a signature specific to the EVs that was not found in the cells. This might be the result of active packaging of some proteins over others in the context of JCPyV infection. Of particular significance was the finding that the proteomic signature of the EV-enriched fraction from JCPyV-infected astrocytes was starkly different from that of the EV-enriched fraction generated under pro-inflammatory cytokine-stimulated conditions. The latter comprised a proteomic signature relating to cytokine-mediated signaling pathways, antigen processing and presentation and type I interferon production. Interestingly, these pathways were not implicated in JCPyV conditions, suggesting that the virus was able to mitigate the cell’s innate immune response as was recently reported for JCPyV in SVG-A cells [41] and BK virus in primary renal proximal tubule epithelial cells [32].

To conclude, while PML has traditionally been characterized as a demyelinating disease associated with the lytic infection of oligodendrocyte, our study, along with those of others [16, 22, 23], emphasizes on the potential role of astrocytes in JCPyV propagation. Although much remains to be done to ascertain the significance of astrocytes in PML onset, we have demonstrated that our model can serve as an excellent tool to address some of the unanswered questions surrounding JCPyV pathophysiology, thereby opening new avenues towards the development of antiviral strategies. We went a step further by demonstrating that EVs secreted from infected astrocytes reflect on-going processes within the cells, thereby indicating the relevance of using astrocyte-derived EVs from body fluids as a window into the CNS and elucidate the obscurity surrounding JCPyV infection in the brain.

## Methods

### Differentiation of human induced pluripotent stem cell (hiPSC)-derived astrocytes

Human iPSCs were generated from peripheral blood mononuclear cells (PBMCs) of two healthy donors (Age/sex: HD2: 50/M; HD3: 49/F) as previously described [17, 42]. The differentiation into astrocytes has been detailed in another publication [43]. Briefly, hiPSCs were first differentiated into neural progenitor cells (NPCs) as per our detailed protocol. Then NPCs were seeded on Matrigel®-coated flasks at 50 000 cells/cm^2^ and kept in Astrocyte Induction medium (DMEM/F-12 supplemented with 1x N2 supplement, 1x B27 supplement, 10ng/ml FGF-2 (PeproTech) and 10ng/ml EGF (Miltenyi)) for two weeks. The cells were thereafter cultured in astrocyte medium supplemented with 20 ng/ml CNTF (PeproTech) for 4 weeks. When mature astrocytes were obtained, the cells were then supplied with Astrocyte medium without CNTF. All donors gave their written informed consent according to institutional review board guidelines. This study was accepted by our institution’s review board (2018_01622). Human iPSC-derived astrocytes were then seeded at [15 000-30 000] cells/cm^2^ a day before the experiment to be performed.

### Infection of hiPSC-derived astrocytes with JCPyV Mad-1

JCPyV Mad-1 (Biogen) was produced as previously described [44]. Human iPSC-derived astrocytes, seeded the previous day, were infected with JCPyV Mad-1 at 8.6 x 10^3^ GE/cell or mock-infected as a control. Following a 24-hour incubation period at 37°C, the cells were washed with 1x phosphate buffered saline (PBS) to remove excess viral particles. Fresh culture media was added to the cells, which were left to incubate at 37°C in a CO2-incubator.

### Stimulation of hiPSC-derived astrocytes with cytokines

A day after seeding, hiPSC-derived astrocytes were supplied with medium containing 10 ng/ml TNFα (R&D Systems) and 10 ng/ml IL-1β (Miltenyi Biotec), or unsupplemented medium as a control. Following a 7-day incubation period, the culture medium of the cells was collected and replaced with fresh TNFα- and IL-1β-supplemented medium for a further 7 days.

### Quantitative PCR (qPCR) analysis

At defined timepoints post-infection (day 3, 7, 14, 21), JCPyV-infected and mock-infected hiPSC-derived astrocytes and supernatants for each corresponding time point were collected and stored at - 20°C. Astrocytes were washed with 1 x PBS and detached using TrypLE (Gibco^TM^). The cells were collected by centrifugation at 600 x g for 10 min and the dry cell pellets were stored at -20°C until further analysis. DNA isolation was performed on the cell pellets and 200 µl of the corresponding supernatants using the QIAamp DNA Blood Mini Kit (Qiagen) according to the manufacturer’s instructions. For the cell pellets, purified DNA samples were standardized according to DNA concentration, as determined by NanoDrop (Thermo Fisher Scientific). A 78 bp amplicon, located on the large T antigen coding region of JCPyV Mad-1, was amplified by specific primers: 5’-AGAGTGTTGGGATCCTGTGTTTT-3’ (nt 4298 to 4320) and 5’-GAGAAGTGGGATGAAGACCTGTTT-3’ (nt 4375 to 4352) and detected with a fluorogenic probe 5’FAM-TCATCACTGGCAAACATTTCTTCATGGC-TAMRA3’ (nt 4323 to 4350) [45].

The reactions were performed in a final volume of 25 µl that consisting of 12.5 μl 2x TaqMan Universal PCR Mix (Applied Biosystem), 400 nM of each primer, 100 nM of the probe and either 10 ng DNA (cell pellet) or 10 μl DNA solution (supernatant). The real-time qPCR reactions were performed using the QuantStudio 6 Flex real-time PCR system (Applied Biosystems), of which the parameters were set to 50°C for 2 min; 95°C for 10 min; followed by 40 cycles of 95°C for 15 sec and 60°C for 1 min. For the preparation of the standard curve, a 10-fold serial dilution in H20 was prepared of a commercially available bacterial plasmid (pBR322) containing the JCPyV Mad-1 full sequence (girt from Peter Howley, Addgene plasmid # 25626) [46]. The final standard curve ranged from 1 to 10^6^ copies/well. For the cell pellets, the results were calculated as copy number per µg total DNA and for the supernatants as copy number per ml culture supernatant. Statistical analysis was performed using GraphPad Prism® 9 software (Version 9.1.0). A Kruskall-Wallis non-parametric repeated measures ANOVA was used to test the overall effect of JCPyV infection overtime. If the effect of the JCPyV infection reached an overall significance, Dunn’s multiple comparisons tests were performed comparing D3 to subsequent timepoints. A p-value of < 0.05 was considered as significant for adjusted P values.

### Immunofluorescence assay (IFA)

At defined timepoints of infection (day 3, 7, 10, 14, 21), JCPyV-infected and mock-infected astrocytes were detached using TrypLE (Gibco^TM^) and replated at 30 000 cells/cm^2^ on 8-well Milicell EZ glass chamber slides (Millipore). Following a 24-hour incubation period, the cells were fixed with 4% paraformaldehyde (PFA) (Electron Microscopy Sciences) and incubated at room temperature for 15 min. The cells were then washed with 1 x PBS and stored in 70% ethanol in H20 at 4°C until further analysis. Once the samples from all the time points were collected, the cells were washed with 1 x PBS and blocked/permeabilized with an IFA solution containing 0.2% Tween-20 (Sigma-Aldrich) and 5% normal goat serum (NGS) (Jackson ImmunoResearch) in 1 x PBS for 1 hour at room temperature. This was followed by incubation at room temperature for 1 hour with the primary antibody in IFA solution (refer to Table 1 for details). The cells were thoroughly washed (2x) with 1 x PBS, followed by incubation for 30 min at room temperature with goat anti-mouse IgG Alexa Fluor 488 (1/200; Invitrogen) or donkey anti-rabbit IgG Alexa Fluor 546 (1/200, Invitrogen) in IFA buffer. The cellular nuclei were then stained with 1.67 µg/ml DAPI (Invitrogen) in IFA buffer. After the final washing step, a coverslip (Vetrini Coprioggetto) was mounted onto the slide using Fluoromount-G (Invitrogen), whereafter the mounted slides were left to dry in a cool, dark place. Cells were analyzed using the LSM 880 confocal with Airyscan microscope (Zeiss) where after images were processed using the Zen (blue edition, v3.6) microscopy software (Zeiss) and quantified using CellProfiler (v4.2.1). A total of 30 images (∼1500 cells), across 3 experiments, were analyzed for each time point and condition using a global thresholding strategy with the Otsu method. Statistical analysis was performed using GraphPad Prism® 9 software (Version 9.1.0). A two-way non-parametric repeated measures ANOVA was used to test the overall effect of JCPyV infection overtime. If the effect of the JCPyV infection reached an overall significance, post hoc Sidak’s multiple comparisons tests were performed at multiple time points. A p-value of < 0.05 was considered as significant for adjusted P values.

### Flow cytometry analysis

JCPyV-infected and mock-infected astrocytes were detached using TrypLE (Gibco^TM^) and labeled with aqua blue amine dye (Life Technologies) for 25 min at 4°C. Cells were washed (2x) with 1 x PBS and stored in 4% PFA (Electron Microscopy Sciences) until analysis with LSRII flow cytometer (BD Biosciences) and FlowJo software (v10.7.1, Treestar). Statistical analysis was performed using GraphPad Prism® 9 software (Version 9.1.0). A two-way non-parametric repeated measures ANOVA was used to test the overall effect of JCPyV infection overtime. If the effect of the JCPyV infection reached an overall significance, post hoc Sidak’s multiple comparisons tests were performed at multiple time points. A p-value of < 0.05 was considered as significant for adjusted P values.

### Fluorescent *in situ* hybridization (FISH)

At defined timepoints of infection (day 3, 7, 10, 14, 21), JCPyV-infected and mock-infected astrocytes were detached using TrypLE (Gibco^TM^) and replated at 30 000 cells/cm^2^ on 8-well Milicell EZ glass chamber slides (Millipore). The next day, cells were fixed with 4% PFA (Electron Microscopy Sciences) in 1 x PBS and stored in 70% ethanol at 4°C. Cells were sequentially dehydrated with 90% and 100% ethanol, for 2 min each, whereafter the hybridization solution, consisting of 1 ug/ml of JC virus BIO-PROBE^®^ (Enzo Life Sciences) in 1x *In Situ* Hybridization buffer (Enzo Life Sciences), was added dropwise to the cells. Coverslips were placed over the slides and sealed with Fixogum (Fisher Scientific), taking care not to form air bubbles. The slides were then placed on a heating block for 5 min at 90°C, to allow DNA denaturation to occur. This was followed by hybridization at 37°C overnight in a humidity-chamber. The next day, the coverslips were removed, and the slides were washed with 1 x PBS, followed by *In Situ* Hybridization Wash Buffer (Enzo Life Sciences) for 15 min at 37°C. The slides were washed with 1 x PBS (2x) and incubated with 1% BSA (Sigma-Aldrich) for 5 min at room temperature. The cells were then incubated with 5 ug/ml Peroxidase Streptavidin (Jackson Immuno Research) in 1% BSA for 30 min at room temperature. The slides were washed again with 1 x PBS (2x), whereafter the revelation solution, consisting of 1x Alexa Fluor 568 Tyramide Reagent (Invitrogen) and 1/2000 dilution of H2O2 (Sigma-Aldrich) in 50 mM Tris buffer at pH 7.4, was added dropwise to the cells and left for 2 min at room temperature. The slides were washed with 1 x PBS (2x) followed by nuclear staining with 1.67 µg/ml DAPI (Invitrogen) in 1 x PBS for 10 min at room temperature. Fluoromount-G (Invitrogen) was used to mount coverslips (Vetrini Coprioggetto) on the slides that were left to dry in a cool, dark place. The cells were analyzed using the DMi1 Inverted microscope (Leica), whereafter the images were processed using the LAS X software (Leica) and quantified using CellProfiler (v4.2.1). A total of 30 images (∼1500 cells), across 3 experiments, were analyzed for each timepoint and condition using a global thresholding strategy with the Otsu method. Graphical representations were performed on GraphPad Prism® 9 software (Version 9.1.0). Statistical analysis was performed using GraphPad Prism® 9 software (Version 9.1.0). A two-way non-parametric repeated measures ANOVA was used to test the overall effect of JCPyV infection overtime. If the effect of the JCPyV infection reached an overall significance, post hoc Sidak’s multiple comparisons tests were performed at multiple time points. A p-value of < 0.05 was considered as significant for adjusted P values.

### Transmission electron microscopy (TEM)

JCPyV-infected and mock-infected astrocytes were fixed with 2.5% glutaraldehyde (Sigma-Aldrich) in 1 x PBS and sent to the Electron Microscopy Facility at the University of Lausanne for sample preparation and TEM analysis. The cells were incubated with a fresh mixture of osmium tetroxide 1% (EMS) and 1.5% of potassium ferrocyanide (Sigma-Aldrich) in 1 x PBS for 1 hour at room temperature. The samples were washed (3x) with distilled H2O and spun down in low melting 2% agarose (Sigma-Aldrich) in H2O and left to solidify on ice. The samples were then cut into 1 mm^3^ cubes and dehydrated in acetone (Sigma-Aldrich) solution at graded concentrations (30% for 40min, 50% for 40min, 70% for 40min and 100% for 1-2h). This was followed by infiltration in Epon (Sigma-Aldrich) at graded concentrations (Epon 1/3 acetone for 2h; Epon 3/1 acetone for 2h, Epon 1/1 for 4h; Epon 1/1 for 12h) and finally polymerized for 48 hours at 60°C in the oven. Ultrathin sections of 50 nm were cut on a Leica Ultracut (Leica) and transferred onto a copper slot grid (2 x 1 mm; EMS) coated with a polystyrene film (Sigma-Aldrich). Sections were post-stained with 2% uranyl acetate (Sigma-Aldrich) in H2O for 10 minutes, followed by rinsing with H2O. The sections were stained with Reynolds’s lead citrate (Sigma-Aldrich) in H2O for 10 minutes and rinsed for a final time with H2O. Micrographs were taken with a transmission electron microscope Philips CM100 (Thermo Fisher Scientific) at an acceleration voltage of 80kV with a TVIPS TemCam-F416 digital camera (TVIPS).

### Analysis of JCPyV-infected astrocytes by liquid chromatography-tandem mass spectrometry LC-MS/MS using a tandem mass tag (TMT)

#### Protein digestion of whole-cell lysate

At different time points of infection, JCPyV-infected and mock-infected cells were lysed with 5 M Guanidine in 50 mM HEPES, pH 8.5. The protein concentrations were determined by using the Pierce^TM^ BCA protein assay kit (Thermo Scientific) and following the manufacturer’s instructions. Aliquots containing 20 μg total protein were prepared and sent to the proteomic core facility (PCF) of EPFL for sample preparation and LC-MS/MS analysis. The samples were digested using the Filter-Aided Sample Preparation (FASP) protocol with minor modifications [47]. Protein samples were resuspended in 8 M urea solution in 100 mM Tris-HCl and deposited on top of Microcon®-30K devices (Merck). Samples were centrifuged at 9400 × g, at 20°C for 30 min. All subsequent centrifugation steps were performed using the same conditions. Reduction was performed using 10 mM Tris(2-carboxy)phosphine (TCEP) in 8 M urea solution. This was followed by the alkylation step using 40 mM chloroacetamide (CAA) in 8 M urea solution and incubation at 37°C for 45 min in the dark. The alkylation solution was removed by centrifugation followed by washing with 8 M urea. Proteolytic digestion was performed overnight at 37°C using a combined solution of Endoproteinase Lys-C and Trypsin Gold in an enzyme/protein ratio of 1:50 (w/w) supplemented with 10 mM CaCl2. The resulting peptides were recovered by centrifugation and desalted on SDB-RPS StageTips and dried by vacuum centrifugation. A mixture of each biological sample was spiked as one channel and used as a bridge channel in all the 3 individual TMT sets.

#### TMT labelling and fractionation

For TMT labelling, dried peptides were first reconstituted in 8 μl of 100 mM HEPES, pH 8 to which 4 μl of TMT solution (25μg/μl pure acetonitrile) was then added. TMT Labelling was performed with the TMT10plex^TM^ isobaric Mass Tagging Kit (Thermo Fisher Scientific) at room temperature for 90 min, whereafter reactions were quenched with hydroxylamine to a final concentration of 0.4% (v/v) for 15 min. TMT-labelled samples were then pooled at a 1:1 ratio across all samples. A single shot LC-MS control run was performed to ensure similar peptide mixing across each TMT channel to avoid the need of further excessive normalization. The quantities of each TMT-labelled sample were adjusted according to the control run. The combined samples were then desalted using a 100 mg Sep-Pak C18 cartridge (Waters) and vacuum centrifuged. Pooled samples were fractionated into 12 fractions using an Agilent OFF-Gel 3100 system following the manufacturer’s instructions. Resulting fractions were dried by vacuum centrifugation and again desalted on SDB-RPS StageTips.

#### LC-MS/MS

Each individual fraction was resuspended in 2% acetonitrile in 0.1% formic acid, whereafter nano-flow separations were performed on a Dionex Ultimate 3000 RSLC nano UPLC system on-line connected with a Lumos Fusion Orbitrap Mass Spectrometer. A capillary pre-column (Acclaim Pepmap C18; 3 μm-100 Å; 2cm x 75 μM ID) was used for sample trapping and cleaning. Analytical separations were performed at 250 nl/min over a 150 min biphasic gradient on a 50 cm long in-house packed capillary column (75 μm ID; ReproSil-Pur C18-AQ 1.9 μm silica beads; Dr. Maisch). Acquisitions were performed through Top Speed Data-Dependent acquisition mode using 3s cycle time. First MS scans were acquired at a resolution of 120’000 (at 200 m/z) and the most intense parent ions were selected and fragmented by High energy Collision Dissociation (HCD) with a Normalised Collision Energy (NCE) of 37.5% using an isolation window of 0.7 m/z. Fragmented ions scans were acquired with a resolution of 50’000 (at 200 m/z) and selected ions were then excluded for the following 120s.

#### Isolation of extracellular vesicles by ultracentrifugation

Extracellular vesicles were isolated from the cell supernatant by differential ultracentrifugation. Briefly, culture medium collected from cells were subjected to centrifugation at 300 x g for 10 min, 2000 x g for 10 min and 10 000 x g for 30 min at 4°C, to clear the conditioned media from cells, cellular debris, and large vesicles, respectively. This was followed by ultracentrifugation at 100 000 x g for 2 hours at 4°C, whereafter the pellet was resuspended in cold 1x PBS and subjected to another round of high-speed centrifugation at 100 000 x g for 2 hours at 4°C. The supernatant was discarded, and the final 100K EV pellet was resuspended in 100 ul cold 1 x PBS. All ultracentrifugation steps were performed using Beckman ultracentrifuge with a SW32Ti rotor. The protein concentrations of the EV isolates were determined using the Pierce^TM^ BCA protein assay kit (Thermo Fisher Scientific) and 10 ug aliquots were prepared and stored at -80°C until further analysis.

### Analysis of EVs by LC-MS/MS using a label-free quantification method (LFQ)

#### Protein digestion

EV aliquots (10 µg) were lysed at a 1:1 ratio with 5 M Guanidine in 50 mM HEPES, pH 8.5 and sent to the proteomic core facility (PCF) of EPFL for sample preparation and LC-MS/MS analysis. Reduction of the samples were done using 10mM TCEP, followed by alkylation of the cysteines using 40 mM CAA for 45 min at 37°C. Samples were diluted with 200 mM Tris-HCl, pH 8, to 1 M guanidine, followed by digestion at room temperature for 2 h with Endoproteinase Lys-C at a 1:50 protease/protein ratio. Samples were further diluted with 200 mM Tris-HCl, pH 8, to 0.5 M guanidine. Trypsin Gold was added at a 1:50 protease/protein ratio, followed by overnight incubation at 37°C. The reaction was quenched with 5% formic acid (FA). Peptides were desalted on SDB-RPS StageTips and dried by vacuum centrifugation.

#### LC-MS/MS

Each individual fraction was resuspended in 2% acetonitrile in 0.1% formic acid and nano-flow separations were performed on a Dionex Ultimate 3000 RSLC nano UPLC system on-line connected with a Lumos Fusion Orbitrap Mass Spectrometer. A capillary pre-column (Acclaim Pepmap C18; 3 μm- 100 Å; 2cm x 75 μM ID) was used for sample trapping and cleaning. Analytical separations were performed at 250 nl/min over a 150 min biphasic gradient on a 50 cm long in-house packed capillary column (75 μm ID; ReproSil-Pur C18-AQ 1.9 μm silica beads; Dr. Maisch). Acquisitions were performed through Top Speed Data-Dependent acquisition mode using 1s cycle time. The first MS scans were acquired at a resolution of 240’000 (at 200 m/z) and the most intense parent ions were selected and fragmented by High energy Collision Dissociation (HCD) with a Normalised Collision Energy (NCE) of 30% using an isolation window of 0.7 m/z. Fragmented ions scans Fragmented ion scans were acquired in the ion trap using a fix maximum injection time of 20ms and selected ions were then excluded for the following 20s.

### Statistical analysis of LC-MS/MS proteomic data

#### TMT data analysis

Raw data were processed using SEQUEST, Mascot, MS Amanda and MS Fragger in Proteome Discoverer v.2.4 against a concatenated database consisting of the Uniprot Homo Sapiens Reference proteome database (75776 protein sequences Release2021_01) and the Uniprot JC Polyomavirus Reference proteome database (8 protein sequences Release2021_01). Enzyme specificity was set to trypsin and a minimum of six amino acids was required for peptide identification. Up to two missed cleavages were allowed and a 1% false discovery rate (FDR) cut-off was applied both at peptide and protein identification levels. For the database search, carbamidomethylation (C), TMT tags (K and peptide N termini) were set as fixed modifications whereas oxidation (M) was considered as a variable one. Resulting text files were processed through in-house written R scripts (v3.6.3). Assuming that the total protein abundances were equal across the TMT channels, the reporter ion intensities of all spectra were summed, and each channel was scaled according to this sum, so that the sum of reporter ion signals per channel equaled the average of the signals across the samples. The multiplexing design of the experiment required a second normalization step to correct variations between the two TMT experiments. Internal Reference Scaling (IRS) process was here applied. A Trimmed M-Mean normalization step was also applied using the package EdgeR (v3.26.8). Differential protein expression analysis was performed using R bioconductor package limma (v3.40.6) followed by the Benjamini-Hochberg procedure. False discovery rate (FDR) < 0.05 were considered as significant.

#### Label-free data analysis

Raw data were processed using MaxQuant (v1.6.10.43) against a concatenated database consisting of the Uniprot Homo Sapiens Reference proteome database (77027 protein sequences Release2021_02) and the Uniprot JC Polyomavirus Reference proteome database (8 protein sequences Release2021_01), Carbamidomethylation was set as fixed modification, whereas oxidation (M), phosphorylation (S, T, Y), acetylation (Protein N-term) and glutamine to pyroglutamate were considered as variable modifications. A maximum of two missed cleavages were allowed and “Match between runs” option was enabled. A minimum of 2 peptides was required for protein identification and the FDR cutoff was set to 0.01 for both peptides and proteins. Label-free quantification and normalization was performed by Maxquant using the MaxLFQ algorithm, with the standard settings. The statistical analyses of the label-free data were performed using Perseus (v1.6.15.0) from the MaxQuant tool suite. Reverse proteins, potential contaminants and proteins only identified by sites were filtered out. Protein groups containing at least 60% or 3 valid values in at least one group were conserved for further analysis. Empty values were imputed with random numbers from a normal distribution. A two-sample t-test with permutation-based FDR statistics (250 permutations, FDR = 0,5; S0 = 1) was performed to determine significant differentially abundant candidates.

#### EV quality check

Rstudio (v4.02) was used to generate a custom R script to annotate proteins considered contaminant proteins versus non-contaminant proteins according to the MISEV2018 specifications [30]. Proteins belonging to the MISEV18 protein content-based EV characterization: 1A, 1B, 2A, 2B, 3A, 3B, were labelled and the sum of the proteins riBAQ in each category was recovered to specify the relative abundance of that category. Note that the proteins categorized with a “*” in the study were manually extended using the Gene Ontology website (Sup. Table. 4). Category 1 included transmembrane or GPI-anchored proteins associated with the plasma membrane and/or endosomes that were either non-cell/tissue specific (1A) or cell/tissue specific (1B). Category 2 comprised cytosolic proteins with lipid or membrane protein-binding ability (2A), and other cytosolic proteins promiscuously associated with EVs (2B). Category 3 served as a purity check that comprised contaminant proteins that might be co-isolated with EV preparations, including lipoproteins and serum-derived materials (3A) or proteins/nucleic acid aggregates and ribosomal proteins (3B). The enrichment of non-contaminant EV enriched categories 1 (A, B) and 2 (A, B) were compared to contaminant categories 3 (A, B) using a nonparametric paired Wilcoxon test.

### Graphical representation

Proteomic data was analyzed using Python (v.3.9.7). Volcano plots were generated using the matplotlib (v3.5.1) for the representation of significantly (FDR ≤ 0.05) dysregulated proteins in the infected or stimulated condition as compared to the mock-infected or resting conditions, respectively. The overlap of significantly dysregulated proteins at different timepoints analyzed were represented by venn diagrams using the venn package (v0.1.3). The proteins were ranked according to the product of |- log10(P) * log2(FC)|, of which the hundred highest-ranked host proteins were used to generate cluster heatmaps. For this, the seaborn package (v0.11.2) was implemented with the seaborn clustermap function that included an average linkage method with a Euclidean metric. The gene list of the hundred highest ranking proteins was subjected for Protein-Protein Interaction (PPI) interaction analysis using the SRTING (v11.5) database with the minimum interaction score set to 0.07 (high confidence). Further analysis of the network was done on Cytoscape (v3.9.1), including Gene Ontology (GO) enrichment analysis using the STRING enrichment plugin. For the category, GO Biological Process was selected with redundancy cut-off of 0.15. All other plots and statistical analyses were done using GraphPad Prism® 9 software (Version 9.1.0).

### Data Availability

Anonymized data not published within this article will be made available by request from any qualified investigator.

## Acknowledgements

We would like to thank Romain Hamelin, Dr. Florence Armand Dr. Maria Pavlou from the Proteomics Core Facility at EPFL for the proteomic analysis, which included all the sample preparations and LC-MS/MS analyses along with the statistical analyses of the different datasets. We would also like to acknowledge and thank Jean Daraspe, Antonio Mucciolo and Dr. Christel Genoud from the electron microscopy facility at the University of Lausanne for with their help with the transmission electron microscopy (TEM) analyses and Dr. Jessica Sordet-Dessimoz from the Histology Core Facility at EPFL for the fluorescent *in situ* hybridization experiments. Many thanks to Dr. Tim Beltraminelli from the De Palma group at EPFL for his insights into characterization of extracellular vesicles (EVs). Confocal images were acquired at the Cellular Imaging Facility at the University of Lausanne.

## Funding sources

This work was supported by the Swiss National Science Foundation (320030_179531), an unrestricted grant from Biogen and a grant from the Myelin Repair Foundation. The funders had no role in study design, data collection and analysis, decision to publish, or preparation of the manuscript.

## Conflict of interests

L.O, A.M., S.P., E.B., M.C., S.J., L.C., M.G., G.C.K., A. S., K. R., S. G. and RDP have nothing to declare related to this work.

## Supporting information

### Supplementary figure legends

**Supplementary** Figure 1**. Transmission electron microscopy (TEM) of mock-infected astrocytes. A, B.** Normal cellular morphologies are represented by two TEM images of mock-infected astrocytes at day 14 post-infection. Mock-infected cells comprised intact plasma membranes and cytoplasms (cyt) with the cell nuclei (nuc) devoid of virus particles and tubular structures.

**Supplementary** Figure 2**. Relative abundance of JCPyV early (LT and ST) and late (VP1 and VP2) proteins over time**. Cells were infected with JCPyV (red) or mock-infected (gray) as described in Fig. 1. At day 0, 3, 7, 14 and 21, the cell lysates were collected and analyzed by LS-MS/MS using a TMT labeling approach. Each dot on the graph represents an experimental replicate and the line the line represents the median. All graphs: n = 3 independent infections performed per readout, with each dot on the graph representing an individual experiment (in red: JCPyV; In grey: mock) and the line links the median value of each condition. The effect of infection over time (D0 vs other timepoints) was tested using a two-way ANOVA followed by Sidak’s multiple comparison test. Statistical significance of data: *p < 0.05; ** p<0.01; ***p < 0.001; ****p < 0.0001.

**Supplementary** Figure 3**. Selection of hundred highest-ranking host proteins in JCPyV-infected conditions.** Cells were infected with JCPyV or mock-infected as described in Fig. 1. Representative scatter plot showing quantified proteins in JCPyV conditions as compared to mock-infected control at day 21 of infection. the Log2(fold-change) of the quantified proteins are represented on the x-axis (JCPyV/ Mock) and the corresponding -Log10(P-value) on the y-axis. Significantly (FDR ≤ 0.05) dysregulated proteins are shown in color, with upregulated proteins shown in red and downregulated proteins shown in blue. The hundred highest-ranked proteins (yellow dots), according to the product of |-log10(P-value) * log2(fold-change)|, were selected for downstream analysis (see Fig. 4)

Supplementary Figure 4. Immunofluorescence analysis (IFA) of DNA damage response (DDR) proteins in mock-infected astrocytes. Representative images taken at 7 d.p.i. of mock-infected astrocytes showing a lack of formation of nuclear foci (yH2AX, BRCA1, FANCD2, p53BP1) and no upregulation of DNA damage checkpoint proteins (ATM, CHEK2, PLK). JCPyV LT is stained in red and VP1 in green (scale bar = 50 µm).

Supplementary Figure 5. Quality check of extracellular vesicles (EVs) isolated from JCPyV-infected or cytokine-stimulated astrocytes. **A, B.** Violin plots representing the abundances (riBAQ score) of EV-associated proteins (categories 1A, 1B, 2A, 2B) as compared to major contaminant proteins typically co-isolated in EV preparations (categories 3A, 3B), according to MISIEV2018 specifications. EV-associated proteins were significantly enriched in all conditions analyzed (**A:** JCPyV vs mock-infected; **B:** cytokine-stimulated vs resting) as compared to contaminant proteins. The significance was determined by using a nonparametric paired Wilcoxon test: * p<0.05; ****, p <0.0001.

**Supplementary** Figure 6**. Selection of the hundred highest-ranked host proteins in EVs from JCPyV-infected or cytokine stimulated conditions.** Human iPSC-derived astrocytes were infected with 8.6 x 10^3^ GE/cell JCPyV or stimulated with 10ng/ml TNFα and 10ng/ml IL-1β. As the negative control for each condition, the cells were either mock-infected or left resting, respectively. **A, B.** Representative scatter plots showing quantified proteins in JCPyV-infected (**A**) or cytokine-stimulated (**B**) conditions at day 14 as compared to mock-infected, or resting controls, respectively. The Log2(fold-change) of the quantified proteins are represented on the x-axis and the corresponding - Log10(P-value) on the y-axis. Significantly (FDR ≤ 0.05) dysregulated proteins are shown in color, with upregulated proteins shown in red and downregulated proteins shown in blue. The hundred highest-ranked proteins in each condition (yellow dots), according to the product of |-log10(P-value) * log2(fold- change)|, were selected for downstream analysis (see Fig. 7)

**Supplementary Table 1.** **Gene Ontology (GO) enrichment analysis of JCPyV-infected astrocytes.** The ten highest-ranked GO terms are indicated with the corresponding FDR value, number of enriched genes and number of background genes in that pathway. The names of the genes are shown in the final column followed by fold-change (underlined) of the gene in the infected conditions as compared to the mock-infected control and corresponding P-value (italic) of the fold-change.

| Term name | Description | FDR | Number of genes | Number of Background genes | Gene names |
| --- | --- | --- | --- | --- | --- |
| GO:0007049 | Cell cycle | 2,40E-68 | 79 | 1313 | RFC2 (<u>1.22</u>, 13.65); KIF22 (<u>1.02</u>, 10.37); TRIP13 (<u>1.07</u>, 11.16); MCM5 (<u>1.66</u>, 12.10); MSH6 (<u>1.22</u>, 13.09); NCAPH (<u>1.19</u>, 8.66); KNSTRN (<u>1.51</u>, 6.90); NCAPG (<u>1.25</u>, 14.05); CCNB1 (<u>2.99</u>, 8.47); KIF23 (<u>1.66</u>, 6.70); KIF11 (<u>2.62</u>, 18.21); NDC80 (<u>1.55</u>, 8.74); MCM4 (<u>1.72</u>, 12.35); PRR11 (<u>2.12</u>, 14.42); LIG1 (<u>1.87</u>, 18.23); MCM6 (<u>1.62</u>, 14.33); MCM2 (<u>1.66</u>, 16.68); ANLN (<u>2.26</u>, 17.92); CDK2 (<u>1.20</u>, 10.29); CCNA2 (<u>1.87</u>, 11.85); SPC25 (<u>1.48</u>, 8.93); CENPH (<u>1.77</u>, 8.45); SMC2 (<u>1.65</u>, 14.63); FANCD2 (<u>1.56</u>, 15.37); CCNB2 (<u>2.22</u>, 8.71); MAD2L1 (<u>1.15</u>, 12.84); PLK1 (<u>1.37</u>, 9.63); TPX2 (<u>1.62</u>, 10.06); RRM1 (<u>1.05</u>, 13.58); BIRC5 (<u>1.12</u>, 9.19); FEN1 (<u>1.27</u>, 12.96); MCM7 (<u>1.73</u>, 14.31); FANCI (<u>1.63</u>, 14.57); ZWILCH (<u>1.04</u>, 9.13); CHAF1B (<u>1.65</u>, 13.47); POLE (<u>1.05</u>, 9.57); KIF15 (<u>1.42</u>, 13.75); NCAPD2 (<u>1.14</u>, 12.11); TACC3 (<u>1.57</u>, 9.95); KNL1 (<u>1.29</u>, 8.60); BLM (<u>2.55</u>, 6.95); UBE2C (<u>1.50</u>, 13.06); SMC4 (<u>1.35</u>, 10.77); RRM2 (<u>1.98</u>, 12.46); IQGAP3 (<u>0.91</u>, 10.64); CENPF (<u>1.45</u>, 11.14); MKI67 (<u>2.26</u>, 15.69); CEP55 (<u>1.55</u>, 7.47); ZWINT (<u>1.65</u>, 9.58); KIF4A (<u>1.40</u>, 11.13); CKS2 (<u>1.99</u>, 10.35); CKAP2 (<u>3.01</u>, 12.07); PCNA (<u>1.03</u>, 12.06); RFC3 (<u>1.43</u>, 14.45); CIT (<u>1.18</u>, 13.07); ECT2 (<u>2.03</u>, 10.58); HELLS (<u>1.44</u>, 12.96); PRC1 (<u>2.43</u>, 8.30); KIF20A (<u>1.43</u>, 8.21); CDK1 (<u>1.93</u>, 14.54); CENPK (<u>1.25</u>, 8.70); WDR62 (<u>1.25</u>, 9.65); NCAPG2 (<u>1.84</u>, 17.78); KIFC1 (<u>2.41</u>, 14.10); TOP2B (<u>0.95</u>, 11.06); RACGAP1 (<u>1.79</u>, 9.03); CDC45 (<u>1.34</u>, 9.37); POLD1 (<u>1.05</u>, 10.70); RFC5 (<u>1.37</u>, 13.78); TOP2A (<u>3.32</u>, 15.44); BRCA1 (<u>1.99</u>, 6.09); PBK (<u>2.55</u>, 15.84); NCAPD3 (<u>0.99</u>, 9.78); TIMELESS (<u>1.11</u>, 8.80); NUSAP1 (<u>2.13</u>, 12.62); KIF18B (<u>2.23</u>, 9.16); UHRF1 (<u>1.03</u>, 12.64); MCM3 (<u>1.61</u>, 12.74); FAM83D (<u>1.43</u>, 7.03) |
| GO:0030261 | Chromosome condensation | 6,11E-13 | 11 | 42 | NCAPH (<u>1.19</u>, 8.66); NCAPG (<u>1.25</u>, 14.05); CCNB1 (<u>2.99</u>, 8.47); SMC2 (<u>1.65</u>, 14.63); NCAPD2 (<u>1.14</u>, 12.11); SMC4 (<u>1.35</u>, 10.77); CDK1 (<u>1.93</u>, 14.54); NCAPG2 (<u>1.84</u>, 17.78); TOP2A (<u>3.32</u>, 15.44); NCAPD3 (<u>0.99</u>, 9.78); NUSAP1 (<u>2.13</u>, 12.62) |
| GO:0000727 | DSB repair via break-induced replication | 6,24E-12 | 8 | 11 | MCM5 (<u>1.66</u>, 12.10); GINS2 (<u>1.48</u>, 11.82); MCM4 (<u>1.72</u>, 12.35); MCM6 (<u>1.62</u>, 14.33); MCM2 (<u>1.66</u>, 16.68); MCM7 (<u>1.73</u>, 14.31); CDC45 (<u>1.34</u>, 9.37); MCM3 (<u>1.61</u>, 12.74) |
| GO:0033045 | Regulation of sister chromatid segregation | 6,11E-09 | 10 | 83 | TRIP13 (<u>1.07</u>, 11.16); CCNB1 (<u>2.99</u>, 8.47); NDC80 (<u>1.55</u>, 8.74); MAD2L1 (<u>1.15</u>, 12.84); PLK1 (<u>1.37</u>, 9.63); BIRC5 (<u>1.12</u>, 9.19); FEN1 (<u>1.27</u>, 12.96); TACC3 (<u>1.57</u>, 9.95); UBE2C (<u>1.50</u>, 13.06); CENPF (<u>1.45</u>, 11.14) |
| GO:0006297 | Nucleotide-excision repair, DNA gap filling | 1,81E-08 | 7 | 23 | RFC2 (<u>1.22</u>, 13.65); LIG1 (<u>1.87</u>, 18.23); POLE (<u>1.05</u>, 9.57); PCNA (<u>1.03</u>, 12.06); RFC3 (<u>1.43</u>, 14.45); POLD1 (<u>1.05</u>, 10.70); RFC5 (<u>1.37</u>, 13.78) |
| GO:0090307 | Mitotic spindle assembly | 2,33E-08 | 8 | 43 | KIF23 (<u>1.66</u>, 6.70); KIF11 (<u>2.62</u>, 18.21); TPX2 (<u>1.62</u>, 10.06); BIRC5 (<u>1.12</u>, 9.19); KIF4A (<u>1.40</u>, 11.13); PRC1 (<u>2.43</u>, 8.30); KIFC1 (<u>2.41</u>, 14.10); RACGAP1 (<u>1.79</u>, 9.03) |
| GO:0051383 | Kinetochore organization | 2,22E-07 | 6 | 18 | NDC80 (<u>1.55</u>, 8.74); CENPH (<u>1.77</u>, 8.45); SMC2 (<u>1.65</u>, 14.63); SMC4 (<u>1.35</u>, 10.77); CENPF (<u>1.45</u>, 11.14); CENPK (<u>1.25</u>, 8.70) |
| GO:0000077 | DNA damage checkpoint | 6,99E-06 | 9 | 140 | CCNB1 (<u>2.99</u>, 8.47); CDK2 (<u>1.20</u>, 10.29); FANCD2 (<u>1.56</u>, 15.37); PLK1 (<u>1.37</u>, 9.63); BLM (<u>2.55</u>, 6.95); DTL (<u>2.16</u>, 11.74); PCNA (<u>1.03</u>, 12.06); CDK1 (<u>1.93</u>, 14.54); BRCA1 (<u>1.99</u>, 6.09) |
| GO:0051988 | Attachment of spindle microtubules to kinetochore | 1,10E-04 | 4 | 13 | KNSTRN (<u>1.51</u>, 6.90); CCNB1 (<u>2.99</u>, 8.47); ECT2 (<u>2.03</u>, 10.58); RACGAP1 (<u>1.79</u>, 9.03) |
| GO:0061982 | Meiosis I cell cycle process | 2,00E-04 | 7 | 114 | TRIP13 (<u>1.07</u>, 11.16); MSH6 (<u>1.22</u>, 13.09); FANCD2 (<u>1.56</u>, 15.37); PLK1 (<u>1.37</u>, 9.63); CKS2 (<u>1.99</u>, 10.35); TOP2B (<u>0.95</u>, 11.06); TOP2A (<u>3.32</u>, 15.44) |

**Supplementary Table 2.**
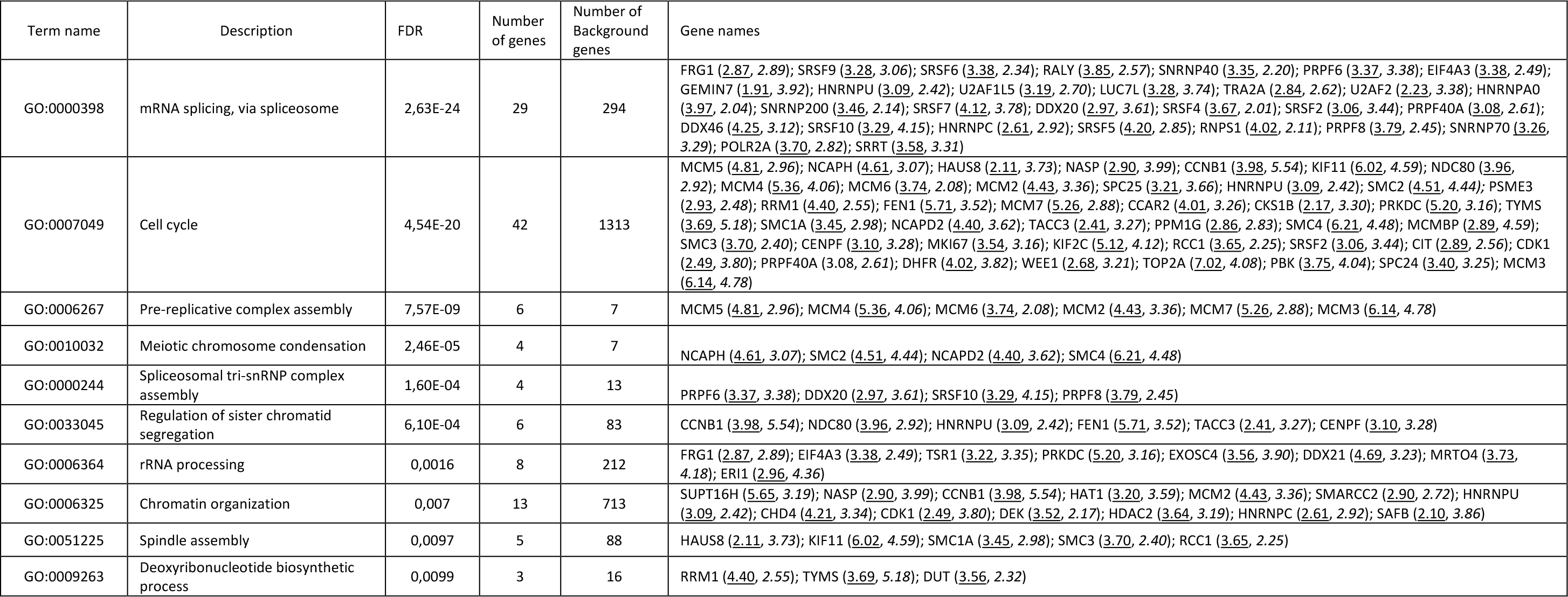
**Gene Ontology (GO) enrichment analysis of EVs from JCPyV-infected astrocytes.** The ten highest-ranked GO terms are indicated with the corresponding FDR value, number of enriched genes and number of background genes in that pathway. The names of the genes are shown in the final column followed by fold-change (underlined) of the gene in the infected conditions as compared to the mock-infected control and corresponding P-value (italic) of the fold-change.

**Supplementary Table 3.**
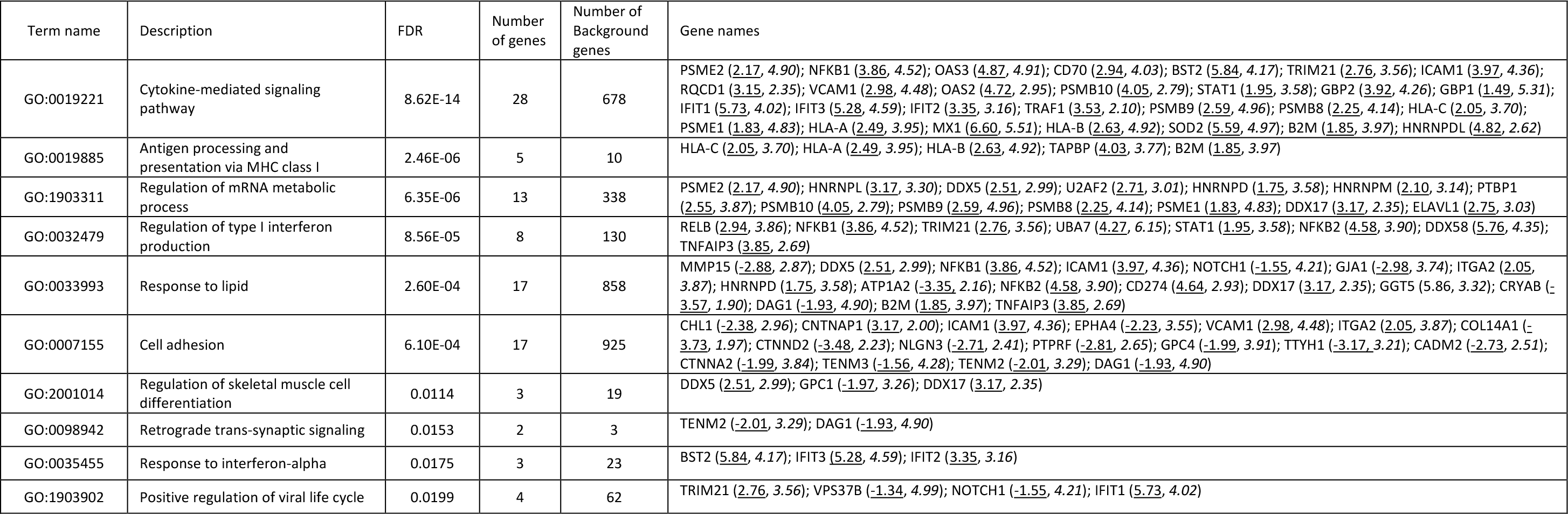
**Gene Ontology (GO) enrichment analysis of EVs from cytokine-stimulated (TNFα+IL-1β) astrocytes.** The ten highest-ranked GO terms are indicated with the corresponding FDR value, number of enriched genes and number of background genes in that pathway. The names of the genes are shown in the final column followed by fold-change (underlined) of the gene in the stimulated conditions as compared to the resting control and corresponding P-value (italic) of the fold-change.

| Term name | Description | FDR | Number of genes | Number of Background genes | Gene names |
| --- | --- | --- | --- | --- | --- |
| GO:0019221 | Cytokine-mediated signaling pathway | 8.62E-14 | 28 | 678 | PSME2 (<u>2.17</u>, 4.90); NFKB1 (<u>3.86</u>, 4.52); OAS3 (<u>4.87</u>, 4.91); CD70 (<u>2.94</u>, 4.03); BST2 (<u>5.84</u>, 4.17); TRIM21 (<u>2.76</u>, 3.56); ICAM1 (<u>3.97</u>, 4.36); RQCD1 (<u>3.15</u>, 2.35); VCAM1 (<u>2.98</u>, 4.48); OAS2 (<u>4.72</u>, 2.95); PSMB10 (<u>4.05</u>, 2.79); STAT1 (<u>1.95</u>, 3.58); GBP2 (<u>3.92</u>, 4.26); GBP1 (<u>1.49</u>, 5.31); IFIT1 (<u>5.73</u>, 4.02); IFIT3 (<u>5.28</u>, 4.59); IFIT2 (<u>3.35</u>, 3.16); TRAF1 (<u>3.53</u>, 2.10); PSMB9 (<u>2.59</u>, 4.96); PSMB8 (<u>2.25</u>, 4.14); HLA-C (<u>2.05</u>, 3.70); PSME1 (<u>1.83</u>, 4.83); HLA-A (<u>2.49</u>, 3.95); MX1 (<u>6.60</u>, 5.51); HLA-B (<u>2.63</u>, 4.92); SOD2 (<u>5.59</u>, 4.97); B2M (<u>1.85</u>, 3.97); HNRNPDL (<u>4.82</u>, 2.62) |
| GO:0019885 | Antigen processing and presentation via MHC class I | 2.46E-06 | 5 | 10 | HLA-C (<u>2.05</u>, 3.70); HLA-A (<u>2.49</u>, 3.95); HLA-B (<u>2.63</u>, 4.92); TAPBP (<u>4.03</u>, 3.77); B2M (<u>1.85</u>, 3.97) |
| GO:1903311 | Regulation of mRNA metabolic process | 6.35E-06 | 13 | 338 | PSME2 (<u>2.17</u>, 4.90); HNRNPL (<u>3.17</u>, 3.30); DDX5 (<u>2.51</u>, 2.99); U2AF2 (<u>2.71</u>, 3.01); HNRNPD (<u>1.75</u>, 3.58); HNRNPM (<u>2.10</u>, 3.14); PTBP1 (<u>2.55</u>, 3.87); PSMB10 (<u>4.05</u>, 2.79); PSMB9 (<u>2.59</u>, 4.96); PSMB8 (<u>2.25</u>, 4.14); PSME1 (<u>1.83</u>, 4.83); DDX17 (<u>3.17</u>, 2.35); ELAVL1 (<u>2.75</u>, 3.03) |
| GO:0032479 | Regulation of type I interferon production | 8.56E-05 | 8 | 130 | RELB (<u>2.94</u>, 3.86); NFKB1 (<u>3.86</u>, 4.52); TRIM21 (<u>2.76</u>, 3.56); UBA7 (<u>4.27</u>, 6.15); STAT1 (<u>1.95</u>, 3.58); NFKB2 (<u>4.58</u>, 3.90); DDX58 (<u>5.76</u>, 4.35); TNFAIP3 (<u>3.85</u>, 2.69) |
| GO:0033993 | Response to lipid | 2.60E-04 | 17 | 858 | MMP15 (<u>-2.88</u>, 2.87); DDX5 (<u>2.51</u>, 2.99); NFKB1 (<u>3.86</u>, 4.52); ICAM1 (<u>3.97</u>, 4.36); NOTCH1 (<u>-1.55</u>, 4.21); GJA1 (<u>-2.98</u>, 3.74); ITGA2 (<u>2.05</u>, 3.87); HNRNPD (<u>1.75</u>, 3.58); ATP1A2 (<u>-3.35</u>, 2.16); NFKB2 (<u>4.58</u>, 3.90); CD274 (<u>4.64</u>, 2.93); DDX17 (<u>3.17</u>, 2.35); GGT5 (5.86, 3.32); CRYAB (<u>-3.57</u>, 1.90); DAG1 (<u>-1.93</u>, 4.90); B2M (<u>1.85</u>, 3.97); TNFAIP3 (<u>3.85</u>, 2.69) |
| GO:0007155 | Cell adhesion | 6.10E-04 | 17 | 925 | CHL1 (<u>-2.38</u>, 2.96); CNTNAP1 (<u>3.17</u>, 2.00); ICAM1 (<u>3.97</u>, 4.36); EPHA4 (<u>-2.23</u>, 3.55); VCAM1 (<u>2.98</u>, 4.48); ITGA2 (<u>2.05</u>, 3.87); COL14A1 (<u>-3.73</u>, 1.97); CTNND2 (<u>-3.48</u>, 2.23); NLGN3 (<u>-2.71</u>, 2.41); PTPRF (<u>-2.81</u>, 2.65); GPC4 (<u>-1.99</u>, 3.91); TTYH1 (<u>-3.17</u>, 3.21); CADM2 (<u>-2.73</u>, 2.51); CTNNA2 (<u>-1.99</u>, 3.84); TENM3 (<u>-1.56</u>, 4.28); TENM2 (<u>-2.01</u>, 3.29); DAG1 (<u>-1.93</u>, 4.90) |
| GO:2001014 | Regulation of skeletal muscle cell differentiation | 0.0114 | 3 | 19 | DDX5 (<u>2.51</u>, 2.99); GPC1 (<u>-1.97</u>, 3.26); DDX17 (<u>3.17</u>, 2.35) |
| GO:0098942 | Retrograde trans-synaptic signaling | 0.0153 | 2 | 3 | TENM2 (<u>-2.01</u>, 3.29); DAG1 (<u>-1.93</u>, 4.90) |
| GO:0035455 | Response to interferon-alpha | 0.0175 | 3 | 23 | BST2 (<u>5.84</u>, 4.17); IFIT3 (<u>5.28</u>, 4.59); IFIT2 (<u>3.35</u>, 3.16) |
| GO:1903902 | Positive regulation of viral life cycle | 0.0199 | 4 | 62 | TRIM21 (<u>2.76</u>, 3.56); VPS37B (<u>-1.34</u>, 4.99); NOTCH1 (<u>-1.55</u>, 4.21); IFIT1 (<u>5.73</u>, 4.02) |

**Supplementary Table 4.**
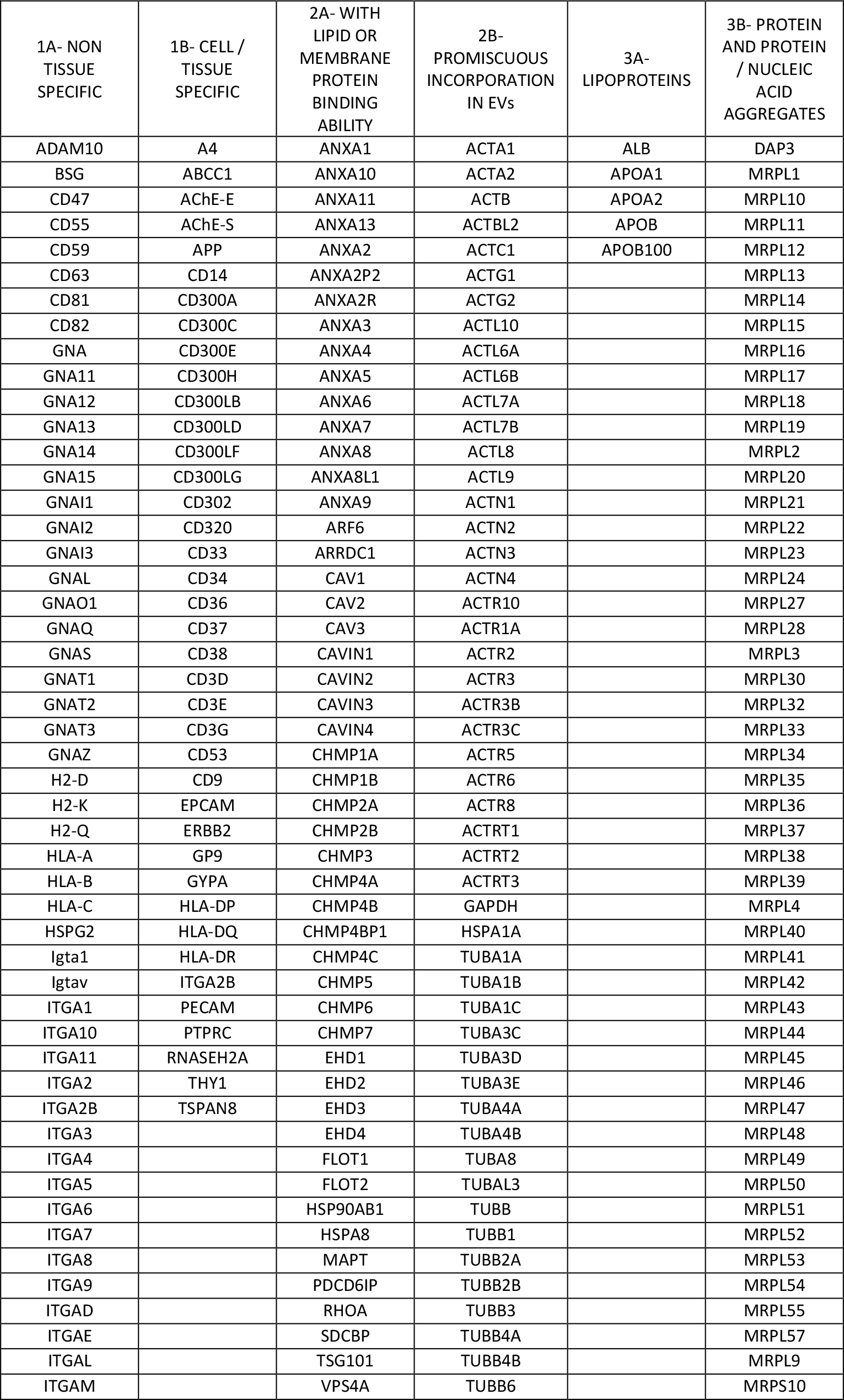
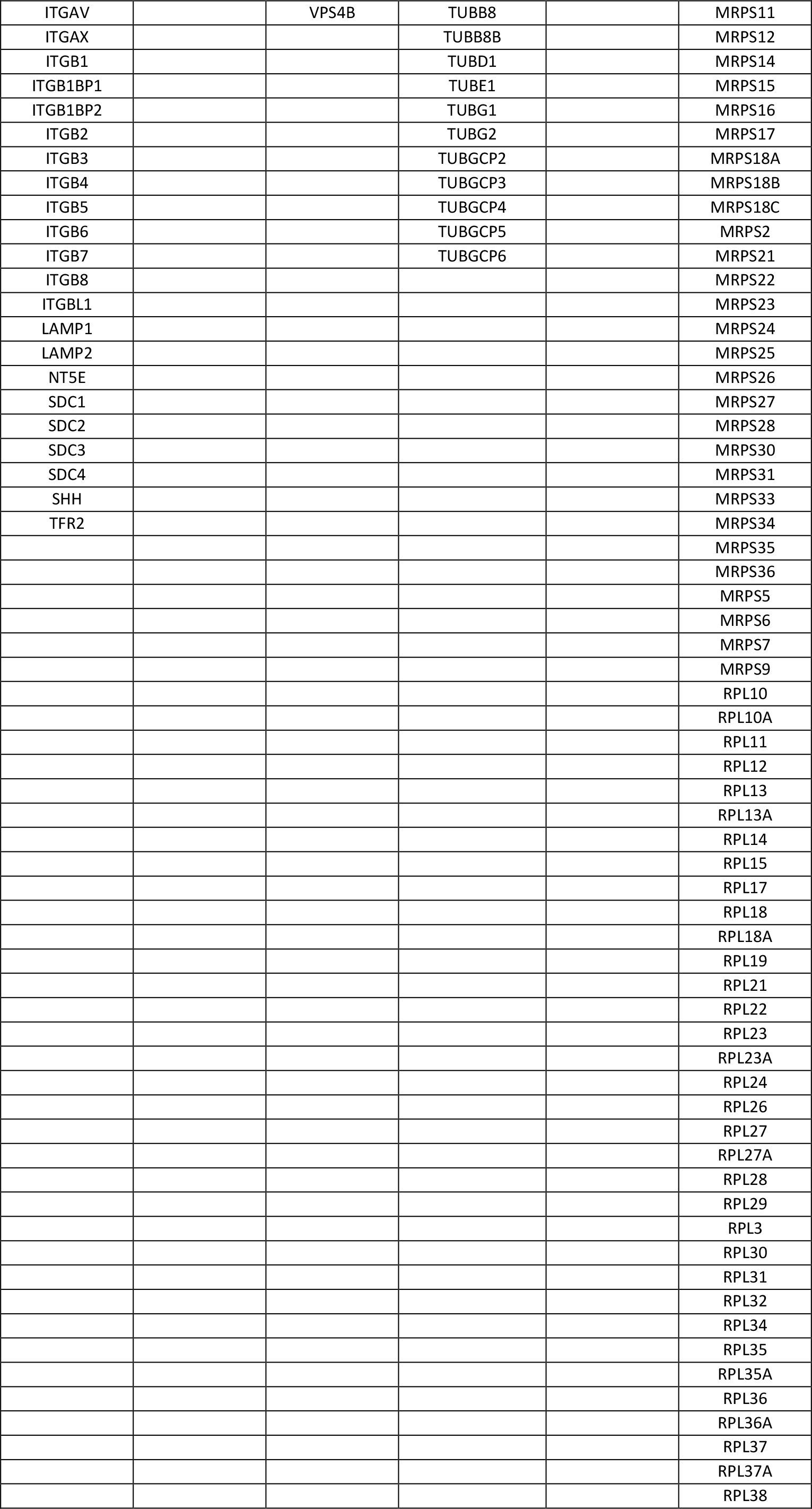
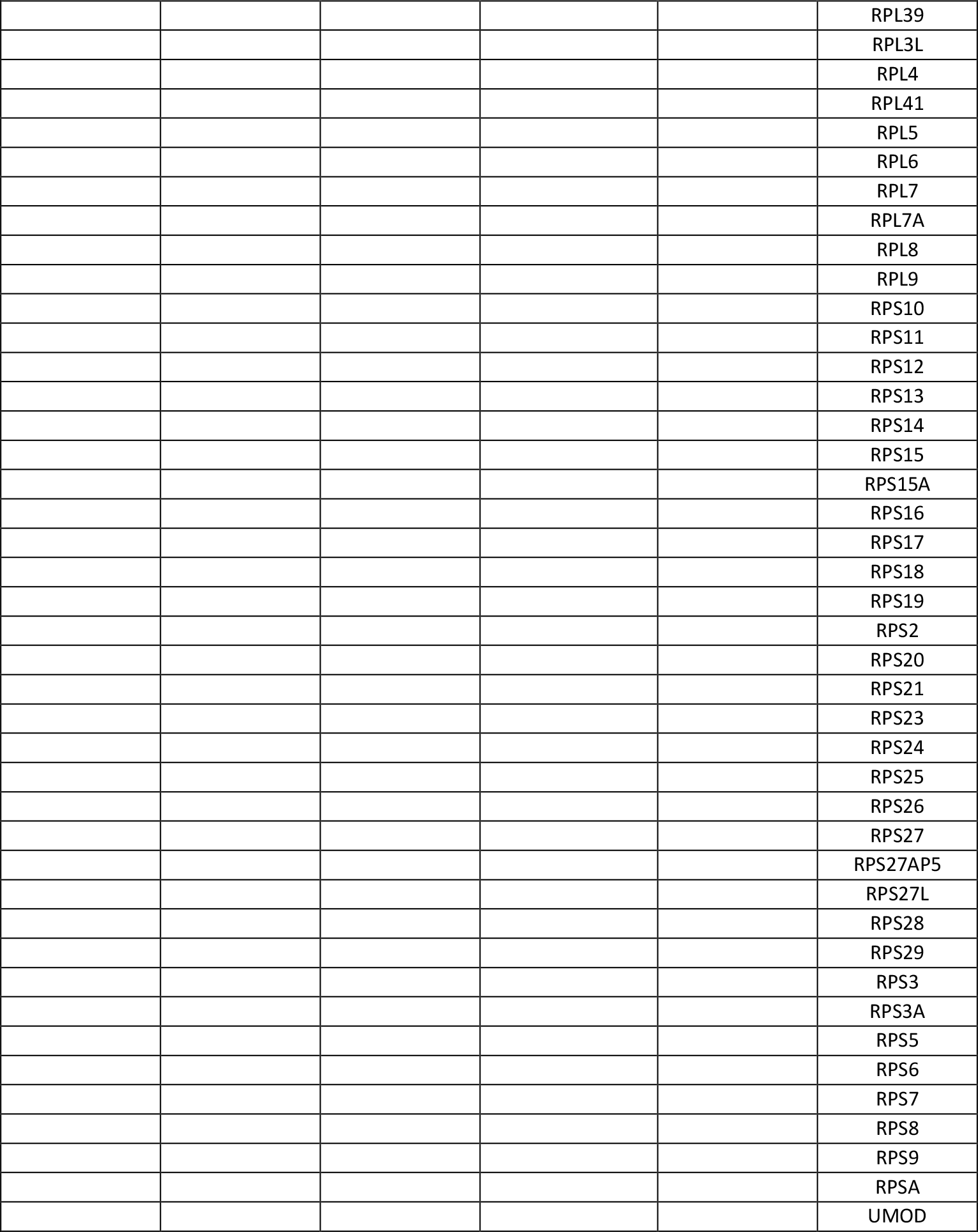
Quality check of EVs based on MISEV2018 specifications [30]. Proteins categorized with a “*” in the study were manually extended using the Gene Ontology website.

## Notes

### Competing Interest Statement

The authors have declared no competing interest.

### Summary of Updates

Minor revisions were made to the abstract, introduction and discussion sections.

