## Supplementary Figure 1 for "Comprehensive proteomic analysis of JC polyomavirus-infected human astrocytes and their excreted vesicles"

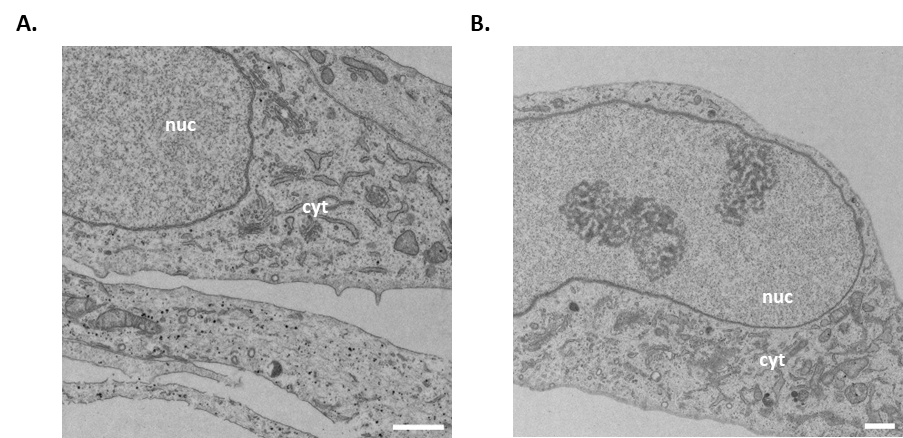

**Supplementary Figure 1. Transmission electron microscopy (TEM) of mock-infected astrocytes.**

**A, B.** Normal cellular morphologies are represented by two TEM images of mock-infected astrocytes at day 14 post-infection. Mock-infected cells comprised intact plasma membranes and cytoplasms (cyt) with the cell nuclei (nuc) devoid of virus particles and tubular structures.
