## Supplementary Figure 2 for "Comprehensive proteomic analysis of JC polyomavirus-infected human astrocytes and their excreted vesicles"

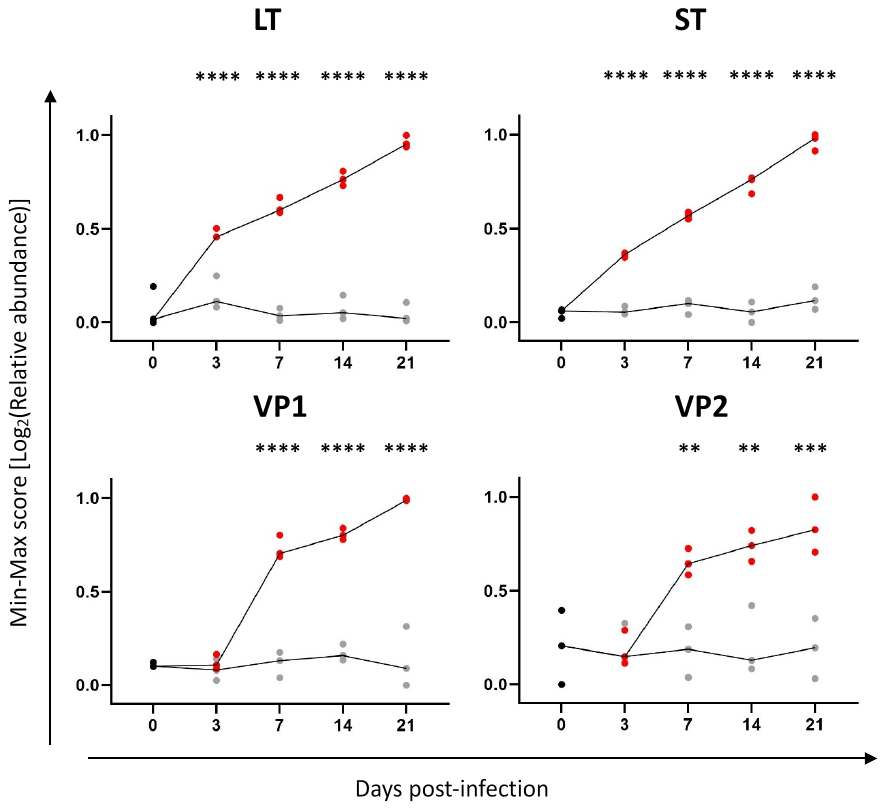

**Supplementary Figure 2. Relative abundance of JCPyV early (LT and ST) and late (VP1 and VP2) proteins over time**. Cells were infected with JCPyV (red) or mock-infected (gray) as described in Fig. 1. At day 0, 3, 7, 14 and 21, the cell lysates were collected and analyzed by LS-MS/MS using a TMT labeling approach. Each dot on the graph represents an experimental replicate and the line the line represents the median. All graphs: n = 3 independent infections performed per readout, with each dot on the graph representing an individual experiment (in red: JCPyV; In grey: mock) and the line links the median value of each condition. The effect of infection over time (D0 vs other timepoints) was tested using a two-way ANOVA followed by Sidak`s multiple comparison test. Statistical significance of data: *p < 0.05; ** p<0.01; ***p < 0.001; ****p < 0.0001.
