## Supplementary Figure 3 for "Comprehensive proteomic analysis of JC polyomavirus-infected human astrocytes and their excreted vesicles"

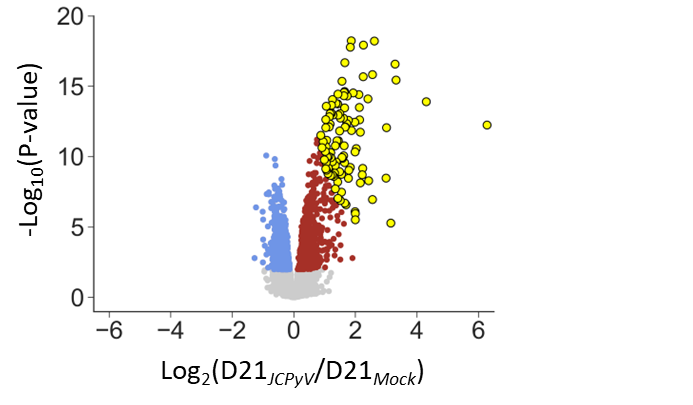

**Supplementary Figure 3. Selection of hundred highest-ranking host proteins in JCPyV-infected conditions.** Cells were infected with JCPyV or mock-infected as described in Fig. 1. Representative scatter plot showing quantified proteins in JCPyV conditions as compared to mock-infected control at day 21 of infection. the Log_2_(fold-change) of the quantified proteins are represented on the x-axis (JCPyV/ Mock) and the corresponding -Log_10_(P-value) on the y-axis. Significantly (FDR ≤ 0.05) dysregulated proteins are shown in color, with upregulated proteins shown in red and downregulated proteins shown in blue. The hundred highest-ranked proteins (yellow dots), according to the product of |-log10(P-value) * log2(fold-change)|, were selected for downstream analysis (see Fig. 4).
