## Supplementary Figure 4 for "Comprehensive proteomic analysis of JC polyomavirus-infected human astrocytes and their excreted vesicles"

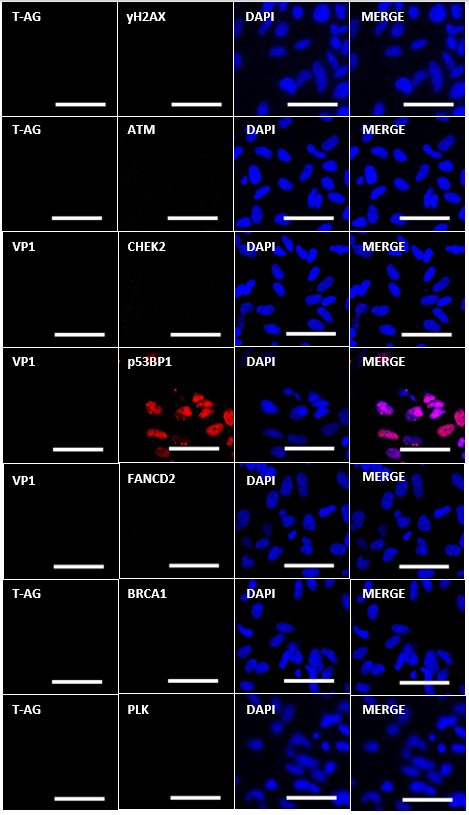

**Supplementary Figure 4. Immunofluorescence analysis (IFA) of DNA damage response (DDR) proteins in mock-infected astrocytes.** Representative images taken at 7 d.p.i. of mock-infected astrocytes showing a lack of formation of nuclear foci (yH2AX, BRCA1, FANCD2, p53BP1) and no upregulation of DNA damage checkpoint proteins (ATM, CHEK2, PLK). JCPyV LT is stained in red and VP1 in green (scale bar = 50 µm).
