## Supplementary Figure 5 for "Comprehensive proteomic analysis of JC polyomavirus-infected human astrocytes and their excreted vesicles"

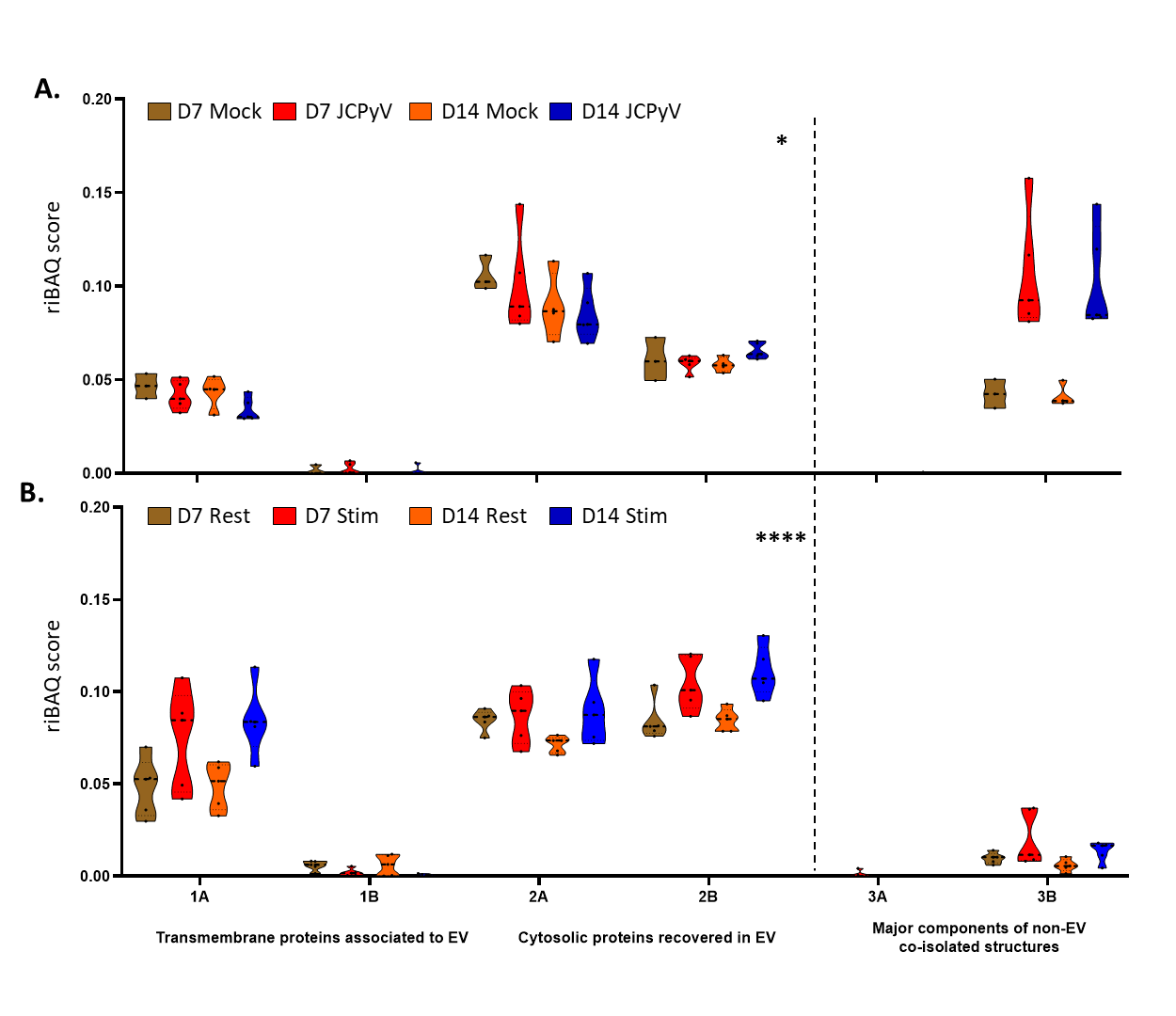

**Supplementary Figure 5. Quality check of extracellular vesicles (EVs) isolated from JCPyV-infected or cytokine-stimulated astrocytes.**

**A, B.** Violin plots representing the abundances (riBAQ score) of EV-associated proteins (categories 1A, 1B, 2A, 2B) as compared to major contaminant proteins typically co-isolated in EV preparations (categories 3A, 3B), according to MISIEV2018 specifications. EV-associated proteins were significantly enriched in all conditions analyzed (**A:** JCPyV vs mock-infected; **B:** cytokine-stimulated vs resting) as compared to contaminant proteins. The significance was determined by using a nonparametric paired Wilcoxon test: * p<0.05; ****, p <0.0001.
