## Supplementary Figure 6 for "Comprehensive proteomic analysis of JC polyomavirus-infected human astrocytes and their excreted vesicles"

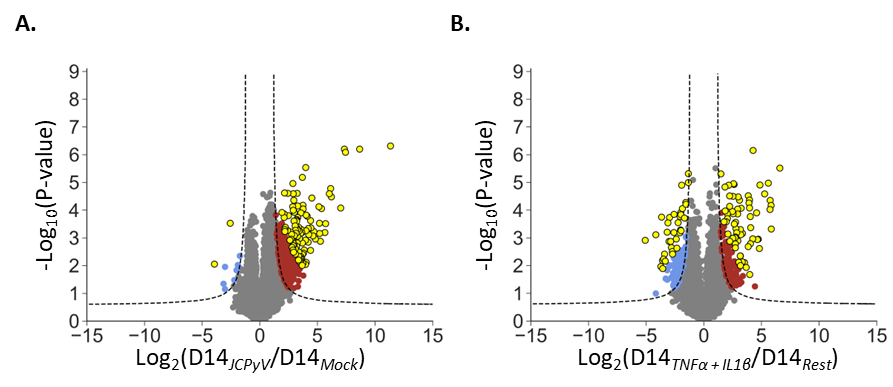

**Supplementary Figure 6. Selection of the hundred highest-ranked host proteins in EVs from JCPyV-infected or cytokine stimulated conditions.** Human iPSC-derived astrocytes were infected with 8.6 x 10^3^ GE/cell JCPyV or stimulated with 10ng/ml TNFα and 10ng/ml IL-1β. As the negative control for each condition, the cells were either mock-infected or left resting, respectively.

**A, B.** Representative scatter plots showing quantified proteins in JCPyV-infected (**A**) or cytokine-stimulated (**B**) conditions at day 14 as compared to mock-infected, or resting controls, respectively. The Log_2_(fold-change) of the quantified proteins are represented on the x-axis and the corresponding -Log_10_(P-value) on the y-axis. Significantly (FDR ≤ 0.05) dysregulated proteins are shown in color, with upregulated proteins shown in red and downregulated proteins shown in blue. The hundred highest-ranked proteins in each condition (yellow dots), according to the product of |-log10(P-value) * log2(fold-change)|, were selected for downstream analysis (see Fig. 7).
